# From hands to tools: evidence for evolutionary precursors of tool processing in macaque cortex

**DOI:** 10.64898/2026.11.24.768489

**Authors:** Yin Liu, Mathilda Froesel, Changshuo Wang, Nora E. Fitzgerald, Gabriel Montaldo, Alan Urban, Ding Cui, Haiyan Wang, Xiaolian Li, Hans Op de Beeck, Stefania Bracci, Rufin Vogels, Qi Zhu, Wim Vanduffel

## Abstract

In humans, hands are animate effectors and tools are visually dissimilar inanimate objects, yet the brain represents them in closely related cortical systems. Whether this coupling depends uniquely on human tool knowledge or reflects a conserved property of primate cortex is unknown. Using whole-brain fMRI (0.22 mm³ voxels) in macaques viewing images of hands, tools, and other objects, we found that inferotemporal cortex largely followed the animate– inanimate division, with only a few small hand-tool preferring sites. In contrast, parietal and prefrontal areas showed hand-tool representational similarities that were best matched by a neural-network model trained on fine-grained human hand-object interactions rather than by networks optimized for static object recognition or general action recognition. Volumetric functional ultrasound imaging (0.0023 mm³ voxels) independently showed that tool responses were spatially aligned with human-hand, but not monkey-hand, responses in posterior parietal cortex. Macaque cortex thus represents tools in relation to the hand even without habitual tool use, consistent with an evolutionarily conserved action-related scaffold that may be shaped by long-term visual exposure to human hand–tool interactions.

## Introduction

Animacy is widely regarded as a fundamental organizing principle of primate high-level visual cortex, evident in both human occipitotemporal cortex and macaque inferior temporal cortex^1–5^. Yet, this broad division does not capture all relationships among categories. For example, hands and tools are highly dissimilar animate and inanimate object categories but are linked by their complementary roles in object-directed action. In humans, static images of hands and tools evoke adjacent, partially overlapping responses in left lateral occipitotemporal cortex, and tool processing engages a left-lateralized network spanning occipitotemporal, parietal, and frontal regions^6–10^ This organization has been attributed to visual exposure to hand–tool interactions, motor experience with tools, and/or conceptual knowledge of tool use^11–13^. Because these forms of experience are tightly coupled in humans, it is unclear whether they are necessary for shared hand–tool representations in the human brain, or instead reflect an evolutionarily conserved visual or visuomotor scaffold shared across primates.

Macaques offer a test. Their visual and visuomotor systems are broadly comparable to ours^14,15^, but they do not spontaneously use tools and lack the culturally accumulated, linguistically mediated tool knowledge characteristic of humans, although some can be trained to use them^16,17^. Observing hand actions engages macaque temporal, parietal (PFG, AIP; see Extended data Table 1 for abbreviations), and premotor (F5) regions that represent the actor, hand–object relations, and action goals^18–23^. These studies demonstrate that hand and object information is integrated during observed actions but do not answer whether this relationship extends beyond the representation of the unfolding action itself. Because dynamic stimuli explicitly specify hand kinematics, object contact, and action goals, the observed integration may depend on online interaction cues rather than reflecting a stable relationship between hands and tools as visual categories. We therefore asked whether static images of hands show a stable relationship with inanimate object categories in macaque cortex when no interaction is depicted, and whether that relationship is specifically strong for tools, as it is in humans.

We addressed this question in two steps. First, we used whole-brain submillimeter-resolution fMRI in awake macaques^24,25^ with the category stimuli previously used in humans^10,11^(Extended data Fig.1). Inferotemporal cortex mainly reflected the animate–inanimate division, whereas dorsal parietal and prefrontal areas showed a closer relationship between hands and tools than between hands and any other category. Artificial neural network comparisons suggested that this geometry reflects learned human hand–object interactions rather than static object or general action recognition.

**Figure 1.**
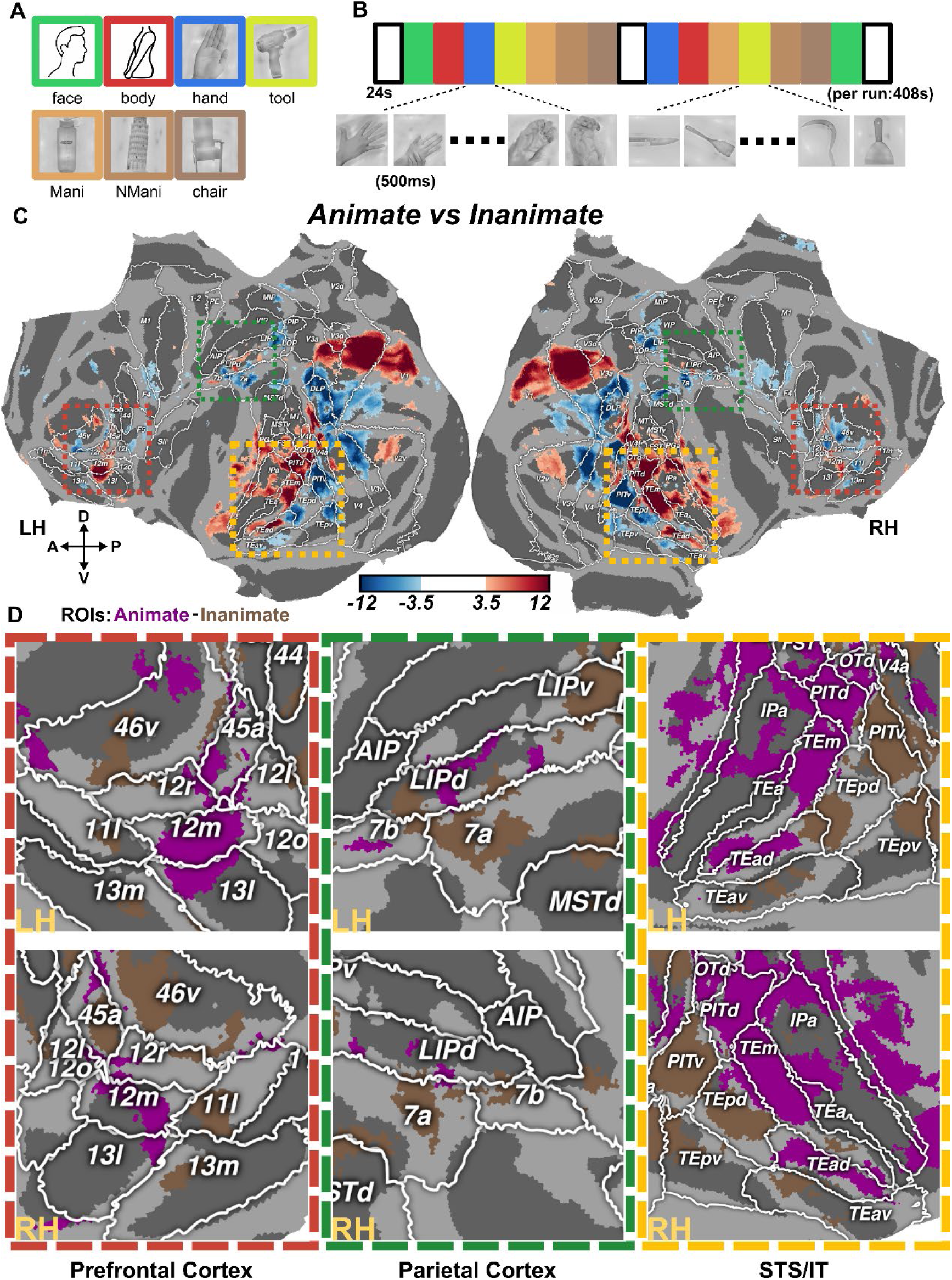
Visual Category Stimuli, Experimental Design, and Whole-Brain Functional Organization of animacy. A. Example stimuli. Representative images from the seven experimental conditions (face, body, hand, tool, manipulable objects, non-manipulable objects, and chair). Human faces and bodies are depicted as icons. The original images of humans used in the experiments are available upon request. B. Experimental design. Each run consisted of 17 blocks (24 s per block), including 14 experimental blocks and 3 baseline blocks (blank screen). Each run began with a baseline block, followed by a first series of seven experimental blocks in which each of the seven conditions was presented once in a randomized order. After an intervening baseline block, a second series of seven experimental blocks was presented in a newly randomized order. The run ended with a final baseline block. Each experimental block contained 48 images (24 original images and their horizontally flipped versions), with each image presented for 500 ms without an inter-stimulus interval. The complete stimulus set is shown in Extended data Fig. 1. C. Group-level cortical surface maps for the animate vs. inanimate contrast Maps are based on the split 1 data across three monkeys. The animate category included faces, bodies, and hands. The inanimate category included tools, manipulable objects, non-manipulable objects, and chairs; Maps show the t-value distribution for the contrast (threshold |t| > 3.5). Warm colors (red) indicate stronger responses to animate stimuli, whereas cool colors (blue) indicate stronger responses to inanimate stimuli. D. Functional regions of interest (fROIs) defined from the animate vs. inanimate contrast. fROIs were defined based on the contrast computed using the split 1 data (p < 0.0001), yielding animate-selective (purple) and inanimate-selective regions (brown). Category assignments were consistent with panel C: animate = face + body + hand; inanimate = tool + manipulable objects + non-manipulable objects + chair.

Second, we predicted that hand-tool similarity would be stronger for human hands than for monkey hands, since our monkeys had never used tools but had seen humans use them on a daily basis. Volumetric functional ultrasound imaging (0.0023mm³ voxels^26^) confirmed this prediction, showing that in parietal cortex tool representations overlapped with those of human but not monkey hands. These findings suggest that tool processing in humans builds on hand-related representations that predate tool use, as they are already present in macaques.

## Results

### Widespread animate–inanimate organization in the macaque cortex

We first characterized the large-scale functional organization of animate and inanimate visual categories across macaque cortex (Fig. 1A, B; Extended data Fig. 1 and 2), both to confirm the previously described animacy-dependent organization of inferotemporal cortex and to ask whether a similar division extends into parietal and prefrontal regions. Whole-brain analyses of 60 of the 121 runs from three monkeys (split 1), with all functional data registered to the MEBRAINS template^27,28^, revealed a widespread and anatomically structured animate–inanimate distinction (Fig. 1C, D). The remaining 61 runs (split 2) were reserved for statistical tests on functional regions of interest (fROI) responses, ensuring independence from fROI definition. Consistent with previous findings in humans and macaques^2,4,5,29,30^, inferotemporal cortex was functionally segregated by animacy. Regions in and surrounding the rostral STS, including IPa, PITd, TEm, and TEad, showed stronger responses to animate stimuli, whereas more caudo-ventral regions, including PITv, TEpd, TEav, and TEpv, showed stronger responses to inanimate stimuli.

Anatomically segregated animate-and inanimate-preferring regions were also identified in parietal and prefrontal cortex (Fig. 1C, D). In prefrontal cortex, however, the two sets of regions were more interdigitated, particularly around areas 46v, 45a,12r/12m, and 13l/13m. A similar organization was observed in parietal cortex, where animate-preferring regions were located predominantly in LIPd and 7b, whereas inanimate-preferring ones were more prominent in area 7a and adjacent LIPv. The animate–inanimate division therefore reaches beyond temporal cortex, extending into parietal and prefrontal regions as well.

### Category-preferring responses show partially overlapping distributions

We next examined how individual object categories contributed to the broad animate-inanimate organization by calculating “category-versus-others” contrasts for faces, bodies, hands, tools, manipulable objects, non-manipulable objects, and chairs (Extended data Fig. 3; Fig. 2). To define the fROIs, we used the split 1 data. A region was termed category-preferring when its response exceeded the mean response of all other categories. All seven categories elicited distributed responses across multiple cortical regions, with both distinct and partially overlapping spatial patterns. For visualization, these maps were displayed at a more liberal threshold (|t| > 2.5) to provide a more inclusive view of the spatial distribution of the responses. As expected^31–33^, face-and body-preferring fROIs were concentrated mainly around the STS and in adjacent inferotemporal areas; additional face-and body-preferring regions appeared in prefrontal and parietal cortex^34^. Hand-preferring fROIs were less prevalent in inferotemporal cortex, appearing instead in parietal and prefrontal regions such as LIPd, area 7b, SII, area 44, and area 12r. Tool and manipulable object preferring responses also extended into parietal and frontal areas, including sectors of area 7b, LIPd, AIP, area 44, and F5. In contrast, non-manipulable objects and chair preferring responses were more prominent in posterior visual and inferotemporal areas, including PITv, TEpd, and TEpv.

### Hand-preferring fROIs range from strict selectivity to shared hand–tool sensitivity

The apparent partial hand-tool co-localization prompted a more focused characterization of hand responses in relation to the other categories. We used the contrasts hand > (face + body) and hand > Inanimate to define relatively strict candidate hand-selective fROIs from their intersection, and broader hand-preferring fROIs from their union (Fig. 3A–B). These fROIs (based on split 1 data) were located in parieto-frontal areas, including 12r and 12m, 44, 46v, F5, SII, 7b, and LIPd, with additional hand-preferring voxels in and around inferotemporal areas TEm and PITd.

We then evaluated response profiles using the split 2 data (Fig. 3C). Only the Intersection fROI in area 7b showed true hand selectivity, with significantly stronger responses to hands than to all six other categories (pairwise t-test, all FDR-corrected *p* < 0.05). In the intersection fROIs of areas 12r, 44, and LIPd, hands elicited the strongest mean response but did not differ significantly from tools. The broader union fROIs showed a similar pattern, with several regions exhibiting comparable responses to hands and tools rather than selective responses to hands only. SII showed relatively strong but more variable responses to both categories, whereas inferotemporal area TEm showed a mixed profile spanning faces, bodies, hands, and tools (Extended data Table 2). The time courses (finite impulse response, FIR, based on split 2 data) were consistent with these activation profiles (Extended data Fig. 4). Hand responses remained distinct in 7b (intersection fROI), whereas hand and tool responses aligned in several other parieto-frontal fROIs. Thus, hand responses in parieto-frontal cortex ranged from relatively strict hand selectivity (only in a subsector of 7b) to shared sensitivity to both hands and tools. The multivariate analyses below directly test representational similarities between hands and tools.

### Spatial overlap of hand and tool preferring regions

hands and tools belong to opposite sides of the animate–inanimate divide and are structurally very different (Extended data Fig. 2), yet their category preferring responses showed similar distributions in dorsal parietal (7b, LIPd) and prefrontal (12r, area 44) cortex (Fig. 2). We quantified this co-localization within atlas-defined cortical areas. For each combination of an animate category (hand, face or body) and an inanimate category (tool, manipulable objects, non-manipulable objects, or chairs), pair-specific localizers were used, with the paired category excluded from the reference condition. Hand–tool overlap, for instance, combined a [hand > others, excluding tool] fROI with a [tool > others, excluding hand] fROI (Extended data Fig. 5). Hand-and tool-preferring responses partially overlapped in several parietal and prefrontal areas, particularly 7b, LIPd, 44, and 12r, and in inferotemporal area TEm (Extended data Fig. 4; Extended data Table 3). These measures are threshold-dependent, and different category pairs can recruit distinct fROIs within the same anatomical area (Extended data Fig. 6). Overlap therefore reflects partial co-localization within an area rather than selectivity of that entire area, so we interpret it only as descriptive evidence.

**Figure 2.**
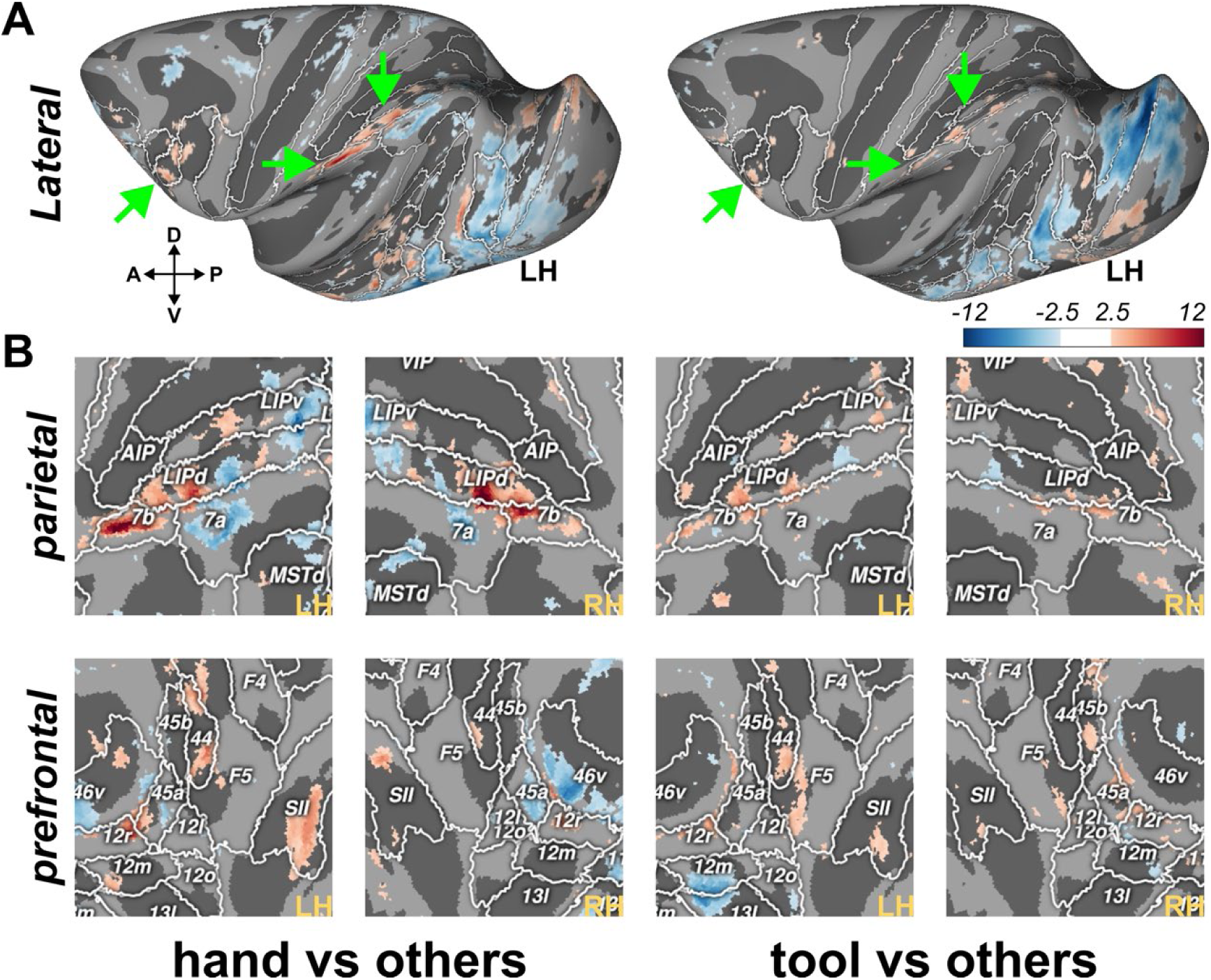
Hand-and tool-preference maps. A. Cortical surface maps for hand-and tool-preference maps. For each contrast, responses to the target category were compared with the average response across the other six categories (using split 1 data). T-score maps are shown on the inflated left hemisphere (dorso-lateral view). Green arrows indicate the prefrontal and parietal regions enlarged in panel B (for both hemispheres). B. Enlarged flattened maps of parietal and prefrontal cortex showing t-scores of the hand-versus-others contrast (left) and tool-versus-others contrast (right), separately for the left (LH) and right (RH) hemispheres. White outlines and labels indicate atlas-defined cortical areas (see Methods: *Region of interest definition*). Warm colors indicate positive t-values and cool colors indicate negative t-values. All maps (A and B) are displayed at ∣ *t* ∣> 2.5.

**Figure 3.**
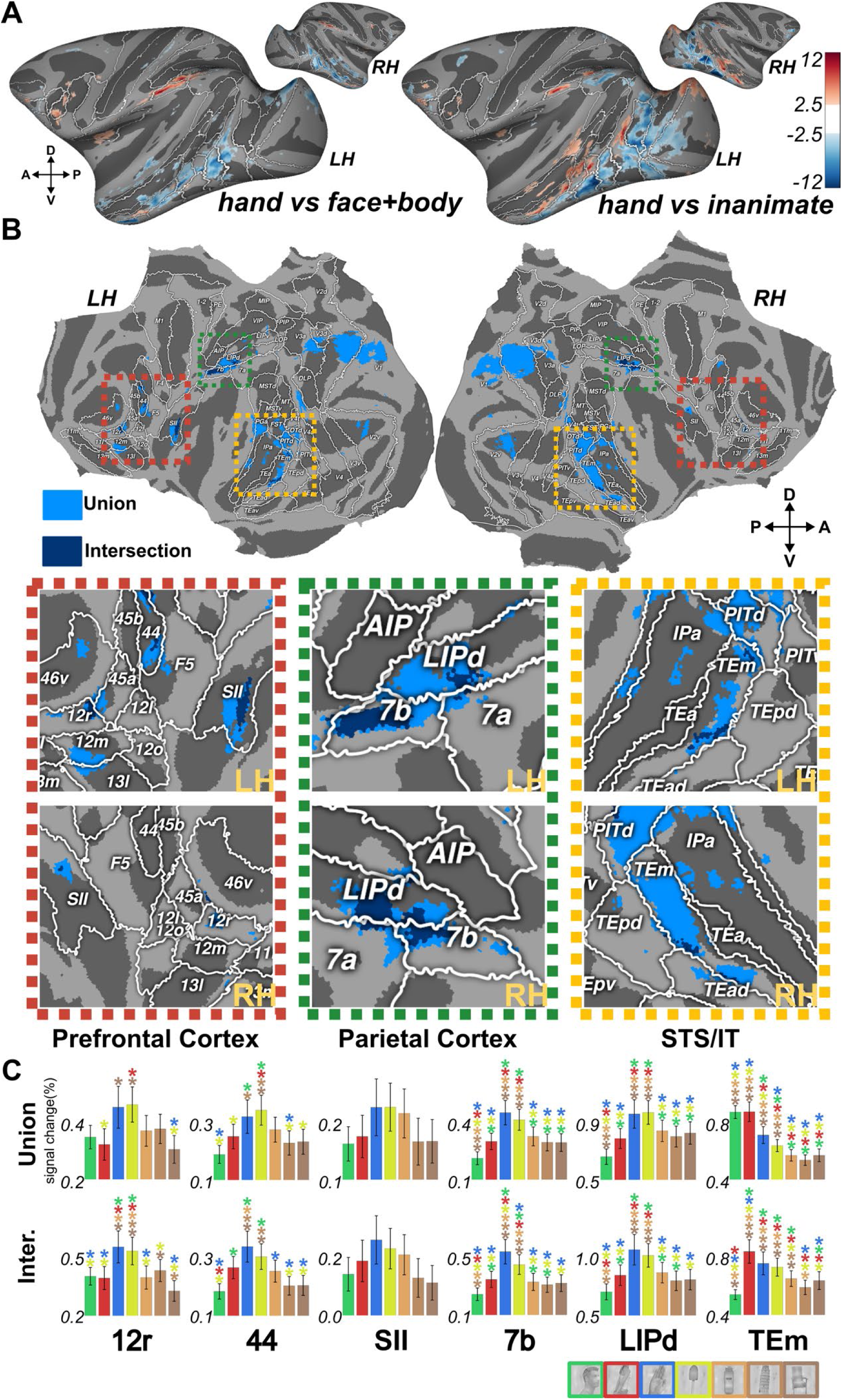
Definition and Validation of hand-Selective Regions. A. Group-level cortical surface maps based on split 1 data for the contrasts hand > (face + body) and hand > Inanimate (tool, manipulable objects, non-manipulable objects, and chair). Maps were thresholded at |t| > 3.5 and used to define hand-selective regions (in C). B. Spatial distribution of hand-fROIs. Two sets of fROIs were defined from the functional localizers (panel A) using the same statistical threshold: “Union” fROIs, comprising the union of the hand > (face + body) and hand > Inanimate contrasts (light blue), and “Intersection” fROIs, comprising their overlap (dark blue). Enlarged views of prefrontal, parietal, and STS/IT cortex illustrate the spatial distribution of the union and intersection hand-fROIs. C. Run-wise activation profiles. Activation profiles (percent fMRI signal change relative to fixation only) were computed from independent split 2 data. Responses are shown for six representative regions (12r, 44, SII, 7b, LIPd, and TEm). Colored asterisks indicate conditions eliciting significantly stronger responses compared to the other categories individually (paired t-test, p < 0.05, FDR-corrected). Only the intersection fROI located within anatomical area 7b shows true hand-selectivity (i.e., hands evoked stronger activity than each of the other conditions). Error bars represent S.E.M. across runs.

### Representational proximity of hands and tools across macaque hand-preferring fROIs

Crossnobis representational dissimilarity matrices (RDMs) were calculated from split 2 data within hand-fROIs independently defined using split 1 (hand vs others excluding tool, *p* < 0.001; Extended data Fig. 7A). Across fROIs, including those overlapping with areas 12r, 44, 7a, 7b, LIPv, LIPd, TEm and PITd, the hand–tool pair generally showed relatively small Crossnobis distances (Fig. 4A). Manipulable objects were also relatively close to hands in some parieto-frontal fROIs, but this pattern was less consistent, whereas non-manipulable objects and chairs did not show a similarly stable relationship with hands across fROIs.

**Figure 4.**
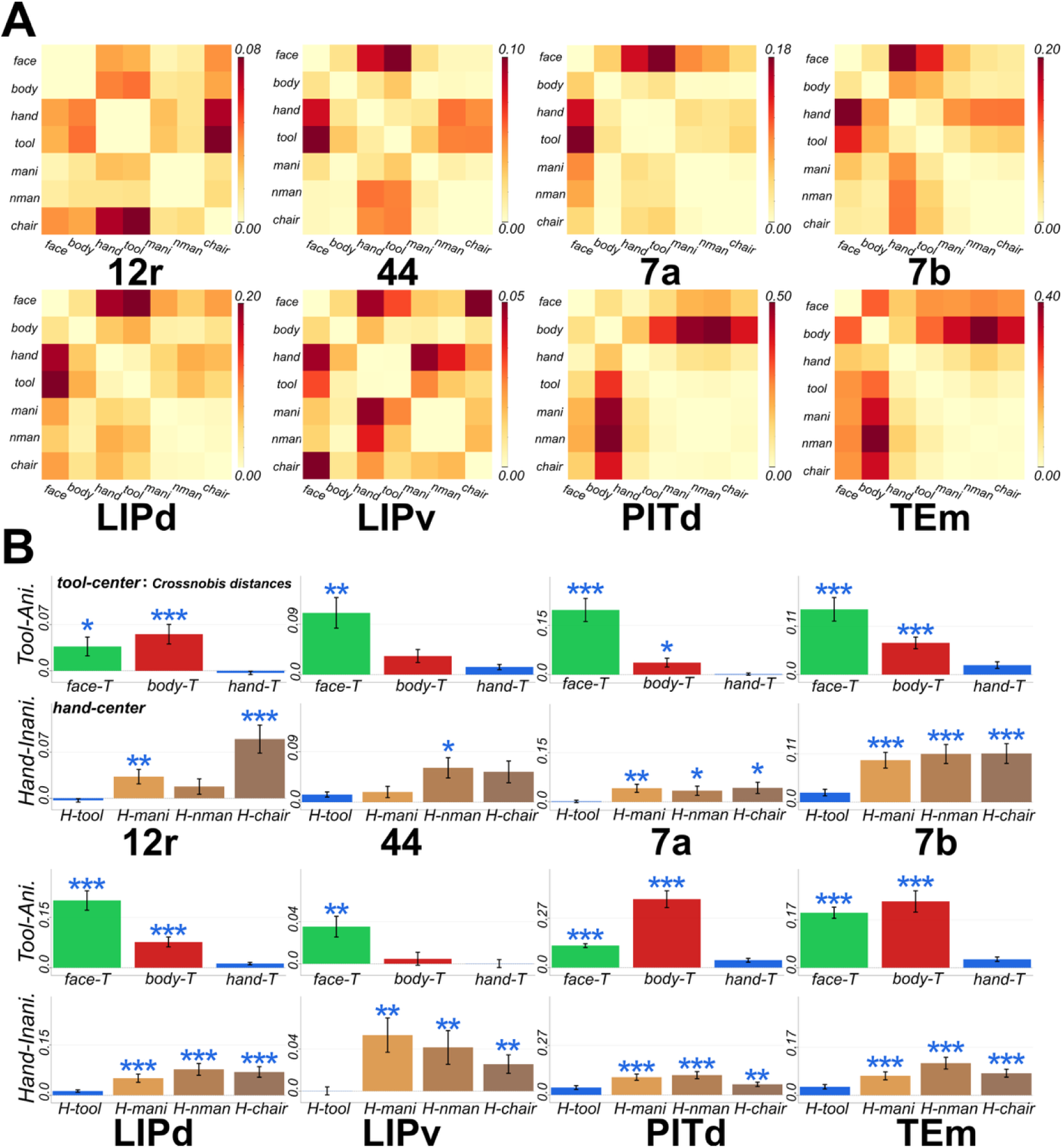
Representational geometry within hand-preferring fROIs. A. Crossnobis representational dissimilarity matrices (RDMs) for hand-preferring fROIs in areas 12r, 44, 7b, 7a, LIPd, LIPv, PITd and TEm. The fROIs were defined on an independent localizer dataset (split 1) using the contrast hand > mean of all other categories except tools (faces, bodies, manipulable objects (mani), non-manipulable objects (nmani) and chairs; P < 0.001). Tools were excluded from the contrast so that the fROI definition was not directly constrained by a potential representational hand-tool relationship. RDMs were computed on the independent test dataset (split 2). Colors indicate the cross-validated Crossnobis distance between each pair of categories, with darker colors denoting greater dissimilarity. B. Tool-and hand-centered Crossnobis distances. For each fROI, the upper panel shows three tool-centered distances (face–tool, body–tool, and hand– tool), whereas the lower panel shows four hand-centered distances (hand–tool, hand–manipulable object [hand–Mani], hand–non-manipulable object [hand– non-manipulable objects], and hand–chair). Hand–tool is displayed in both panels as the shared reference. Bar colors denote the category pairs: face–tool (green), body–tool (red), hand–tool (yellow), hand– manipulable objects (orange), hand– non-manipulable objects (light brown), and hand–chair (dark brown), respectively. Bars show the mean cross-validated Crossnobis distance across runs, and error bars indicate the S.E.M across runs. Smaller values indicate greater representational proximity, and values near or below zero indicate no reliably measurable separation. Face–tool, body–tool, hand–Mani, hand– non-manipulable objects, and hand–chair distances were each compared with the hand–tool distance using two-sided paired t-tests across matched runs. The five resulting P values were corrected within each fROI using the Benjamini–Hochberg false discovery rate procedure. *, **, and *** indicate FDR-adjusted P < 0.05, P < 0.01, and P < 0.001, respectively.

We then compared the hand–tool Crossnobis distance with five related distances: face–tool, body–tool, hand–manipulable objects, hand–non-manipulable objects, and hand–chair (Fig. 4B, Extended data Fig. 7B). In 7a, 7b, and LIPd, the hand–tool distance was significantly smaller than all five comparison distances (all FDR-adjusted P < 0.05), indicating a highly consistent hand–tool representational relationship across these parietal fROIs. A similarly robust pattern was observed in the inferotemporal fROIs PITd and TEm, where the hand–tool distance was also significantly smaller than all five comparison distances. The pattern was less uniform in LIPv and the prefrontal fROIs. In LIPv, the hand–tool distance was significantly smaller than the face– tool, hand–manipulable objects, hand–non-manipulable objects, and hand– chair distances, but not the body–tool distance. In 12r, significant differences were observed relative to face–tool, body–tool, hand–manipulable objects, and hand–chair, but not hand–non-manipulable objects. In area 44, significant differences were confined to the face–tool and hand–non-manipulable-object distances. Thus, hand–tool proximity was broadly and consistently expressed across parietal areas 7a, 7b, and LIPd, and also extended to small hand-preferring inferotemporal fROIs PITd and TEm, with more variable effects in LIPv, 12r, and 44. The broader pattern of pairwise distances across all hand-fROIs is summarized in Extended data Fig. 7B. Together, the spatial-overlap and representational similarity analyses provide convergent evidence for a hand–tool relationship in parieto-frontal cortex, while also revealing representational proximity between hands and tools in small inferotemporal fROIs.

### Hand action-oriented models capture the dorsal hand–tool geometry

Having established the spatial and representational relationships among stimulus categories, we next asked which types of representational structure best corresponded to the neural RDMs. We compared four candidate models that together covered static object representations at two hierarchical levels and video-trained action representations learned from two different datasets (Fig. 5A). To benchmark generic visual and category structure, we used the V1 (CORnet-V1) and IT (CORnet-IT) stages of CORnet-S^35,36^, a shallow recurrent network trained on ImageNet whose processing areas mirror macaque ventral visual areas. To test whether action-oriented training changes this geometry, we used two Video Swin Transformer models with an identical architecture that differed only in their training data. One was trained on Kinetics-400 (K400)^37^, a broad human action dataset whose categories include both human–object and human–human interactions. The second model started from the K400-trained weights and was further trained on Something-Something V2 (SSV2)^38,39^, which requires discriminating fine-grained differences in how hands manipulate everyday objects. Comparing SSV2 with K400 therefore isolates the contribution of interaction-focused training rather than that of model architecture. Because both neural and model RDMs were based on static images, this comparison concerns representational geometry rather than motion encoding.

**Figure 5.**
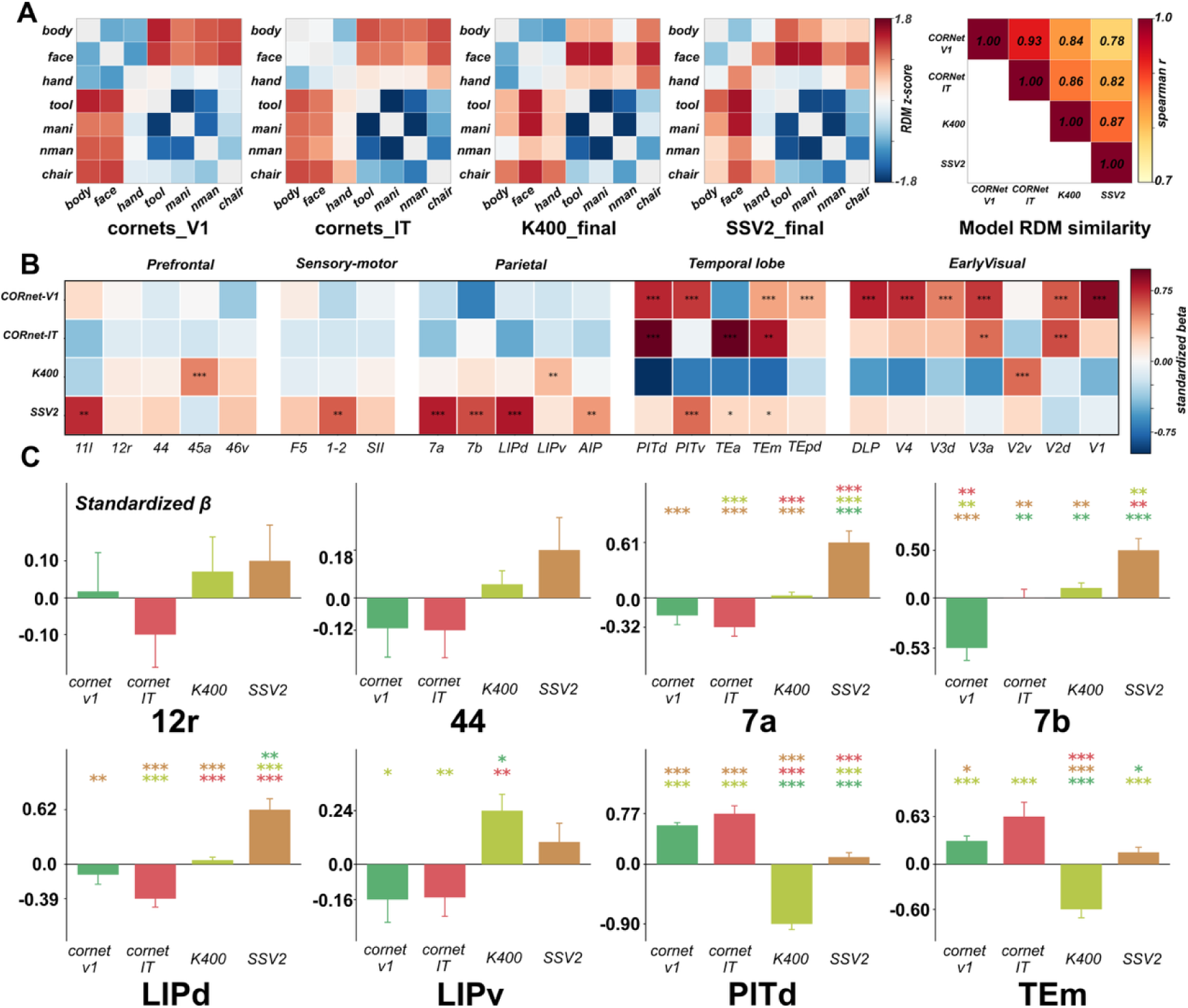
Visual and action-trained models show distinct correspondences with neural representational geometries. A. Model representational dissimilarity matrices (RDMs) and inter-model similarity. RDMs are shown for the V1 and IT stages of CORnet-S (CORnet-V1 and CORnet-IT) and Video Swin Transformer models trained on Kinetics-400 (K400) and Something-Something V2 (SSV2). Each RDM represents seven stimulus categories: bodies, faces, hands, tools, manipulable objects (Mani), non-manipulable objects (NMan), and chairs. For visualization, dissimilarities were z-scored within each model across the 21 unique off-diagonal category pairs and displayed on a common color scale. Red and blue indicate dissimilarities above and below the within-model mean, respectively; diagonal entries were excluded and are shown in gray. The right matrix shows pairwise Spearman rank correlations (ρ) between model RDMs across the same 21 category pairs, with the color scale spanning 0.7–1.0. B. Multiple-regression RSA across hand-preferring fROIs. The four predictor RDMs were entered simultaneously into a multiple linear regression model to explain the neural RDM of each fROI. Hand-preferring fROIs were defined on the independent localizer dataset (split 1) using the contrast hand > all other categories excluding tools, at P < 0.001. Colors indicate the standardized regression coefficient (standardized β) of each predictor, reflecting its unique contribution after accounting for the other three models. Positive coefficients indicate a positive independent correspondence between the model and neural representational geometries. Coefficients were tested against zero using one-sided sign-flip permutation tests across runs, and the resulting P values were corrected across all model-fROI combinations using the Benjamini-Hochberg false discovery rate (FDR) procedure. *, **, and *** indicate FDR-adjusted P < 0.05, P < 0.01, and P < 0.001, respectively. C. Model contributions in representative hand-preferring fROIs. Bars show the mean standardized regression coefficients for CORnet-V1 (green), CORnet-IT (red), K400 (yellow), and SSV2 (brown) in six representative fROIs of areas 12r, 44, 7b, 7a, LIPd, LIPv, PITd, and TEm. Error bars represent the S.E.M. across runs. Pairwise differences between model coefficients were assessed using paired t-tests across runs, and the resulting P values were corrected for the six model comparisons within each fROI using the Benjamini-Hochberg false discovery rate (FDR) procedure. Colored asterisks above each bar indicate a significant difference from the model whose color matches the asterisk. *, **, and *** indicate FDR-adjusted P < 0.05, P < 0.01, and P < 0.001, respectively.

Spearman representational similarity analysis (RSA) showed that CORnet-V1 corresponded mainly to early visual fROIs, whereas CORnet-IT showed stronger correspondence with inferotemporal fROIs (Extended data Fig. 8). K400 was positively related to some early visual and temporal fROIs but showed more limited correspondence in dorsal regions. SSV2 showed broader correspondence, including several parietal and prefrontal fROIs. Multiple-regression RSA further separated the contributions of the models (Fig. 5B, Extended data Table 4). CORnet-V1 retained its clearest positive contribution in early visual fROIs, whereas CORnet-IT contributed more strongly in inferotemporal fROIs. The independent contribution of K400 was substantially reduced after controlling the other models, whereas SSV2 retained a positive independent contribution in several inferotemporal, parietal and prefrontal fROIs. Thus, among the candidate models examined here, the category geometry learned by the hand action-oriented SSV2 model showed the closest correspondence with that of several parieto-frontal fROIs.

The model RDMs were highly correlated (pairwise Spearman ρ = 0.78–0.93). Variance inflation factors (VIFs)^40^ ranged from 6.40 to 11.71 and were highest for CORnet-IT (Extended data Fig. 9A). To assess sensitivity to predictor inclusion, we compared SSV2 coefficients from the full four-predictor model with those obtained after separately omitting CORnet-V1, CORnet-IT, or K400 ^41^, using identical neural data and condition pairs. Across these specifications, mean SSV2 coefficients remained positive in parietal fROIs 7a, 7b, LIPd, and LIPv, with smaller positive means also observed in prefrontal fROIs 12r and 44 (Extended data Fig. 9B).

### 3D-fUSI reveals stronger spatial alignment of tool responses with human than monkey hands in posterior parietal cortex

To test whether visual experience shapes the hand–tool relationship found with whole-brain fMRI, we used 3D-fUSI in posterior parietal cortex of two additional monkeys that had not taken part in those experiments. 3D-fUSI provided substantially higher spatial resolution (0.0023 mm³ voxels within a field of view of approximately 1 cm³) and greater sensitivity than fMRI. In the 3D-fUSI experiment, we divided each of the face, body, and hand stimulus categories into human and monkey categories (Extended Data Fig. 10).This split was motivated by the possibility that macaques, despite lacking direct tool-use experience, have substantial daily visual exposure to human hand–tool interactions but not to monkey hand-tool interactions. If visual exposure contributes to this organization, tool representations should preferentially overlap with those of human rather than monkey hands.

A graded pairwise conjunction analysis revealed category-dependent response patterns across the sampled field of view (Fig. 6). Importantly, the gradation score reflects relative categorical preference rather than response magnitude. A striking spatial correspondence was observed between human-hand and tool-preferring voxels. In both monkeys, several human-hand-preferring clusters were located within, or immediately adjacent to, tool-preferring clusters in and around LIP and area 7a, as well as in the posterior part of the sampled field near the dorsal prelunate area (DP). In contrast, this spatial organization was markedly weaker for monkey hands, for which the corresponding hand - responsive clusters were sparser and largely absent from or spatially distinct from tool-responsive areas. Thus, the local spatial relationship between tools and hands differed across the two hand-image categories, with tool responses aligning more strongly with human-hand than with monkey-hand representations.

**Figure 6.**
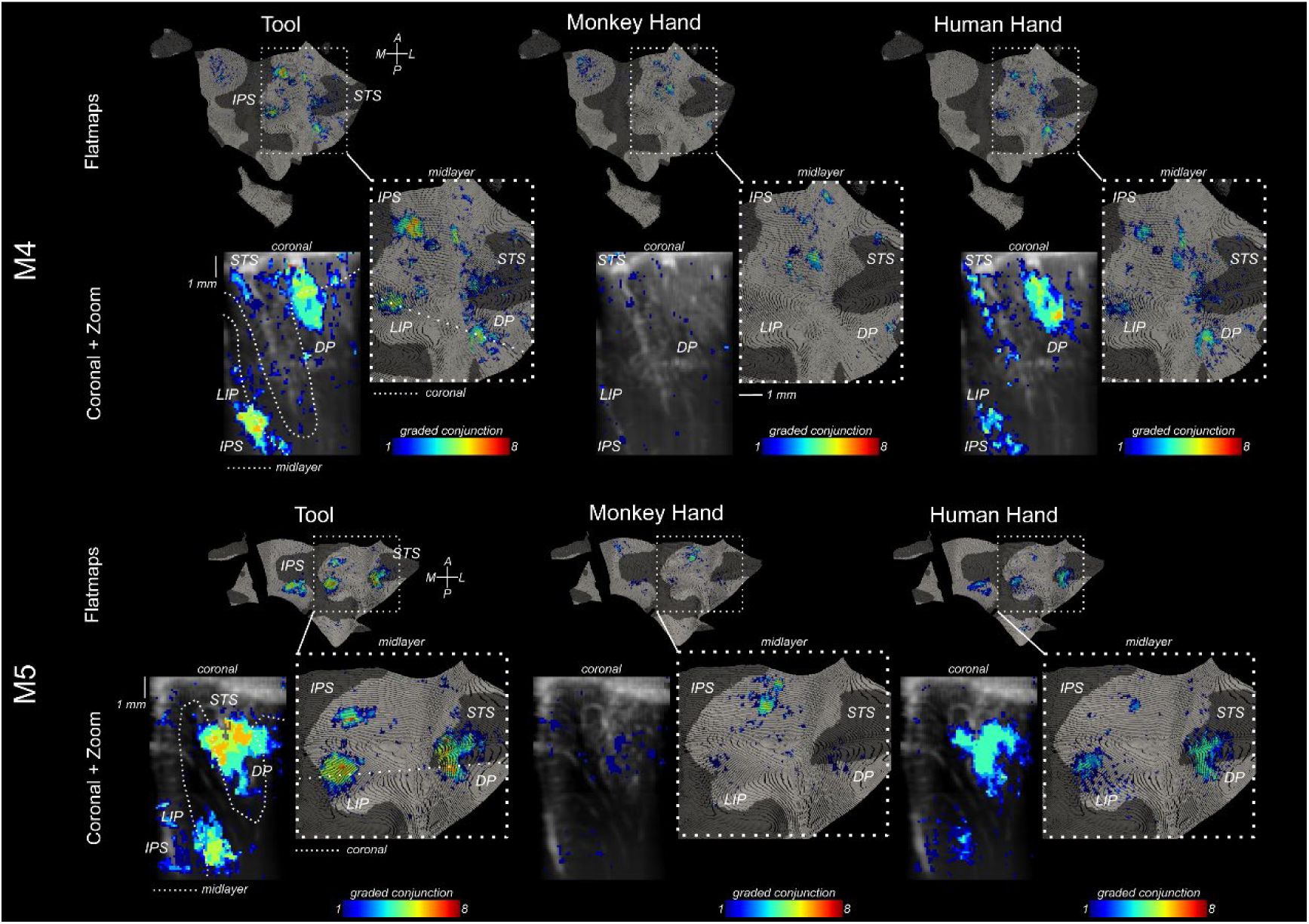
Graded categorical selectivity for tools, monkey hands, and human hands measured with volumetric functional Ultrasound Imaging (3D-fUSI). Graded categorical selectivity for tools, monkey hands, and human hands measured with volumetric functional Ultrasound Imaging (3D-fUSI). Voxel-wise categorical selectivity maps are shown for two monkeys (M4, top; M5, bottom) for tool, monkey hand, and human hand categories. The maps were derived from a graded pairwise conjunction analysis. For each target category, the corresponding fUSI response was independently compared with each of the other eight visual categories (using a general linear model, GLM). (Z > 3.09; p < 0.001, uncorrected). For each voxel (0.0023 mm³), the number of significant pairwise comparisons favoring the target category was summed to give a selectivity score ranging from 1 to 8. Dark blue voxels therefore responded significantly more strongly to the target category than to one other category, and progressively warmer colors indicate increasing categorical selectivity. Voxels with a score of 8 responded more strongly to the target category than to each of the eight other categories separately and therefore correspond to strictly category selective (winner-take-all) voxels. For each monkey and condition, selectivity maps are displayed on the flattened cortical reconstruction (flatmap) of the middle layer of the cortical ribbon (upper panels; each individual dot corresponds to an individual fUSI voxel). Below the flatmap for each selectivity map, representative coronal sections are shown (lower panel, left) as well as a zoomed-in section of the flatmap corresponding to the cortical region between the lateral bank of the IPS and the STS (lower panel, right). In each coronal slice, the dotted line represents the approximate position of the flattened midlayer, similarly, in the zoomed-in flatmap, the dotted line shows the approximate position of its matching coronal slice. Anatomical landmarks and cortical regions are indicated: intraparietal sulcus (IPS), superior temporal sulcus (STS), lateral intraparietal area (LIP), dorsal prelunate area (DP). The scale bars: ∼ 1 mm.

Together, these 3D-fUSI results provide independent fine-scale spatial evidence complementing the parietal hand–tool relationship observed with whole-brain fMRI. The stronger alignment between tools and human hands is consistent with the hypothesis that visual exposure to human hand–object interactions contributes to this organization.

## Discussion

The present study demonstrates that a broad animate–inanimate organization coexists with a more specific hand–tool representational structure in the macaque cortex. Inferotemporal fROIs primarily reflected the global animate– inanimate division, except for small subdivisions of TEm and PITd, whereas parieto-frontal fROIs expressed a closer relationship between hands and tools. This pattern was not explained simply by strict hand selectivity, since only one fROI showed relatively selective responses to hands (a sector of area 7b) while several dorsal fROIs exhibited shared sensitivity to hands and tools. Spatial overlap and representational similarity analyses converged in identifying tools as the inanimate category most consistently related to hands, with manipulable objects showing a weaker and more fROI-dependent relationship. Together, these fMRI findings reveal a regionally specific hand–tool organization that cannot be reduced to broad animacy or to hand selectivity alone. 3D-fUSI in two additional macaques complemented these findings, showing that in sampled parietal cortex tool responses aligned more with human-hand than with monkey-hand responses (Figs. 6, 7B).

**Figure 7.**
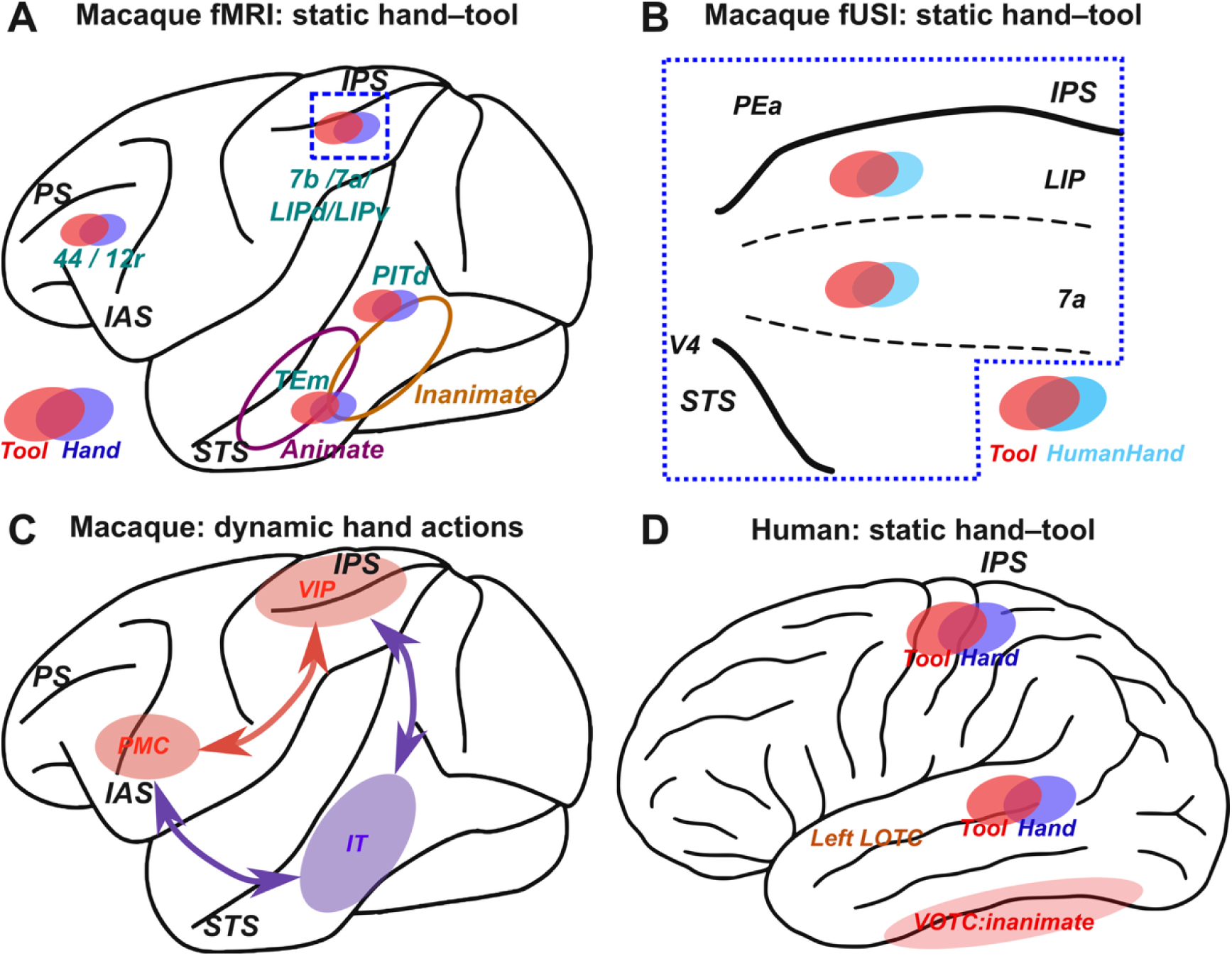
Schematic comparison of static hand–tool organization and dynamic hand-action networks in macaque and human cortex. A. Summary of the macaque fMRI findings. Hand-preferring fROIs were defined using the contrast of hands versus all other categories except tools. In parietal (7a/7b/LIPd/LIPv) and prefrontal (44/12r) cortex, these hand-preferring fROIs partially overlapped with tool-preferring responses. Within the hand-preferring fROIs, representational analyses further showed that, among the tested inanimate categories, tool representations were most consistently closest to hands. B. Summary of the macaque 3D-fUSI findings in a restricted parietal field-of-view. In the 3D-fUSI experiment we presented human-hand and monkey-hand stimuli as distinct categories and evaluated graded pairwise category preferences. In both monkeys, tool-preferring and human-hand-preferring clusters partially overlapped or were spatially adjacent within the sampled field, including clusters around LIP and area 7a. Monkey-hand-preferring responses were surprisingly sparse or absent at these corresponding locations and are therefore not depicted. Blue dotted outlines indicate the fUSI field-of-view. C. Literature-derived summary of the distributed macaque network engaged during the observation of dynamic hand–object actions. Visual analysis of observed actions in and around the STS interacts with parietal and ventral premotor components involved in representing hand–object relations and action goals. Arrows indicate schematic interactions among temporal, parietal, and premotor components but do not represent connections or causal directions tested in the present study^21^. (IAS, inferior arcuate sulcus; IPS, intraparietal sulcus; IT, inferotemporal cortex; PMC, premotor cortex; PS, principal sulcus; STS, superior temporal sulcus; VIP, ventral intraparietal area.) D. Literature-derived summary of human cortical organization during static hand viewing. Hand-and tool-related responses are adjacent or partially overlapping in left lateral occipitotemporal cortex (LOTC) and intraparietal sulcus (IPS), whereas ventral occipitotemporal cortex (VOTC) more strongly retains the broader animate–inanimate organization^10,11,42^. Colored fields indicate qualitative locations and representational relationships; their spatial extent and degree of overlap are schematic and not drawn to scale.

In humans, static images of hands and tools evoke closely adjacent and partially overlapping responses in left lateral occipitotemporal cortex, while tool processing also recruits a broader left-lateralized network spanning parietal and frontal regions (Fig. 7D)^9,10,43,44^. In the present macaque data, convergent spatial and representational relationships between hands and tools were particularly evident in parietal fROIs, including 7a,7b, LIPv, and LIPd, whereas the effects in prefrontal fROIs such as 12r and 44 were more variable (Fig. 7A). At a finer spatial scale, the fUSI results further revealed that tool-preferring responses in macaque parietal cortex were more closely aligned with human-hand than with monkey-hand representations (Fig. 6; Fig. 7B). Importantly, the multivariate analyses also revealed small hand–tool distances in the hand-preferring inferotemporal fROIs PITd and TEm (Fig. 4A,B). This inferotemporal representational proximity did not correspond to a comparably prominent or uniquely identifiable hand–tool territory in the univariate response maps. This proximity instead remained confined to these small fROIs, set against the dominant animate–inanimate organization of inferotemporal cortex. Thus, the macaque results do not support a strict dorsal–temporal dissociation. Hand– tool relationships were present at the representational level in both dorsal and selected inferotemporal hand-fROIs, though they differed across regions in their spatial expression and consistency. Because the two hemispheres were combined in the present analyses, hemispheric lateralization was not directly assessed.

These differences in cortical expression may partly reflect the different forms of experience and knowledge available to each species. Macaques can acquire stable visual and event relationships through repeated observation of hand– object interactions and through their broader experience with grasping and object-directed actions. Human tool experience additionally involves extensive direct manipulation, knowledge of object function, culturally transmitted practices, and linguistic and conceptual information^45,46^. Moreover, tool-selective responses in left LOTC can emerge in the absence of visual experience, indicating that human tool-related organization cannot be explained solely by visual appearance or visual co-occurrence^47^.

The present findings therefore support neither a simple presence–absence distinction between species nor a strictly hierarchical interpretation. Instead, they are consistent with the possibility that macaques and humans share a basic primate visual representational architecture linking hands and tools, while differing in its cortical distribution and degree of specialization. In humans, extensive tool use, functional knowledge, and conceptual experience may further elaborate this organization. This interpretation also resonates with the broader idea of “cultural recycling of cortical maps”^48^, whereby experience-dependent processes shape and repurpose evolutionarily older neural architectures to support species-specific cognitive abilities. However, because the present study did not directly manipulate visual or motor experience, the experiential origins of these differences remain to be established.

Macaque fMRI studies of action observation suggest one route through which experience could shape such an architecture. Observing goal-directed hand actions recruits a distributed network spanning temporal, parietal, and frontal cortex (Fig. 7C)^22^. At the neuronal level, individual cells in the superior temporal sulcus respond both to articulated body actions and to the static postures produced by those actions, suggesting that static configurations can carry information about dynamic events^48,49^. The parieto-frontal distribution of the hand–tool relationship observed here is therefore broadly compatible with the macaque dynamic action observation systems. Static hands and tools may engage representations shaped by prior experience with dynamic hand–object interactions, although we did not test whether static and dynamic stimuli evoke the same representations.

This raises a question about where such an organization comes from. If experience with dynamic hand-object interactions shapes how static categories relate to one another, models trained on those interactions should reproduce dorsal geometry more closely than models optimized for static object identity alone. Our computational comparisons addressed this directly. Representations from the V1 and IT stages of CORnet-S were more closely associated with early visual and inferotemporal fROIs, respectively, as expected for a network designed to approximate the primate ventral hierarchy and trained on static objects^35,36^. SSV2 by contrast retained a more pronounced unique association with several parietal and prefrontal fROIs.

Although the two Video Swin models shared one architecture, their training data imposed very different classification demands. Kinetics-400 spans broad action categories for which scene context, characteristic objects, body posture, and overall appearance often provide sufficient cues for recognition. By contrast, SSV2 forces discrimination among fine-grained human hand–object interactions that differ in hand–object relations, temporal order, movement direction, or changes in object state^37–39^. Because the two networks are identical apart from this training difference, and because both were evaluated on static images here, the stronger SSV2 correspondence points to a specific conclusion. Interaction-focused training can reorganize how static inputs are represented, so images of hands, tools, and manipulable objects come to retain relationships derived from their typical roles in dynamic events.

This does not mean dorsal fROIs directly encode motion, since neither our neural nor our model analyses used moving stimuli. More generally, RSA establishes similarity between representational geometries rather than equivalence of the underlying computations. Our results therefore support neither a strict division in which temporal cortex represents categories and dorsal cortex represents actions, nor an exclusive action-based reading of the SSV2 effect. Shape, posture, affordance, object state, familiarity, and dataset-specific visual statistics could each contribute to what we observed. What the comparison does show is that training data can shift representational geometry in ways relevant to how hands relate to tools, given that K400 and SSV2 differ only in this respect. This in turn raises the question whether a comparable structure could arise in the brain through mere exposure to such interactions. Although the monkeys had not received systematic tool-use training, their long-term experience in a human-managed environment provided repeated opportunities to observe hand–tool interactions. Such exposure may foster associations through spatial proximity, temporal continuity, and recurring action sequences. This possibility is consistent with evidence that associative learning can bring neuronal representations of paired stimuli closer together and that temporal experience can reshape object representations in the macaque visual system^50–53^. The hand–tool proximity observed here may therefore reflect, at least in part, representational structure accumulated through repeated exposure to hand–object interactions.

Consistent with this account, 3D-fUSI revealed stronger spatial alignment of tool responses with human than with monkey hand responses in the sampled parietal cortex (Fig. 6). These findings are compatible with experience-dependent shaping of hand–tool representations, building on pre-existing neural substrates and motor or somatosensory experience. The monkeys’ greater visual exposure to human hand–tool interactions may have been a key driver of this process, although its causal contribution remains to be established. Macaque cortex represents hands and tools in relation to one another even without habitual tool use. Strict hand selectivity was rare, found in only one fROI (within area 7b), while inferotemporal cortex largely followed the animate– inanimate division. Several parietal and prefrontal fROIs nonetheless showed hand–tool similarity, and this similarity matched networks trained on human-object interactions rather than static object recognition. Functional ultrasound independently showed that, in sampled parietal cortex, tool responses aligned more closely with human-hand than with monkey-hand responses. These results indicate a regionally specific hand-tool organization that does not require uniquely human tool knowledge and is consistent with an evolutionarily conserved action-related scaffold that may be shaped by long-term visual exposure to human hand-tool interactions. In humans, this scaffold may then be elaborated by tool use and conceptual knowledge.

## Materials and methods

### Subjects and general procedures

Five rhesus macaques (*Macaca mulatta*) participated in the study. Three of them (M1 (monkey D), M2 (monkey F), M3 (monkey T); two males and one female; approximately 7, 9, and 6 kg, respectively) participated in the fMRI experiment. Two male monkeys (M4 (monkey G), M5 (monkey D)) took part in the fUSI experiment, weighing approximately 7.5 and 8.5 kg, 12 and 11 years of age. All animals were implanted with an MR-compatible headpost for head stabilization during experiments^54^. For the fMRI animals, a multi-channel phased-array receive coil was embedded in the headpost above the skull to increase sensitivity for submillimeter-resolution imaging^24,25,55^.

All monkeys were trained to maintain fixation on a central fixation point while seated in a primate chair with the head immobilized by the headpost. Eye position was monitored continuously, and correct fixation behavior was reinforced with liquid reward. For the fMRI experiment, fixation was required to remain within a 2° × 2° virtual window. To minimize susceptibility artifacts caused by body motion, the monkeys were additionally trained to keep both hands inside a small response box positioned in front of the chair. Hand position was continuously monitored using two pairs of fMRI-compatible optical fibers, and rewards were delivered only when both fixation and hand-position criteria were satisfied^24^.

Animal care and experimental procedures were performed in accordance with the National Institutes of Health *Guide for the Care and Use of Laboratory Animals* and European legislation (Directive 2010/63/EU) and were approved by the Animal Ethics Committee of KU Leuven. Animal housing and handling followed the Weatherall report guidelines. Animals were group-housed in enclosures of approximately 16–32 m³ and provided with environmental enrichment, including foraging devices and toys. They received standard primate chow supplemented with fruits, vegetables, bread, peanuts, cashew nuts, raisins, and dried apricots. Animals participating in the experiments were maintained under a controlled water-access protocol and could work until satiated; on non-working days, water was provided in their home enclosure.

### fMRI acquisition

Functional MRI data were acquired while the monkeys were awake using a 3 T Siemens PrismaFit scanner at KU Leuven. Whole-brain functional images were collected using a two-dimensional simultaneous-multi slice gradient-echo echo-planar imaging sequence at 0.6-mm isotropic resolution. The repetition time was 3,000 ms, and the echo time was 22 ms for M1 and 21 ms for M2 and M3. Images were acquired with a multiband factor of 2, an in-plane acceleration factor of 3, partial Fourier acquisition of 5/8, and a flip angle of 90°. The in-plane field of view was 84 × 84 mm with a matrix size of 140 × 140. Seventy-four coronal slices were acquired for M1 and M2, whereas 90 slices were acquired for M3 to accommodate differences in individual brain coverage. Custom receive coils with 8, 10, and 16 active channels were used for M1, M2, and M3, respectively, together with a custom-built local single-loop transmit coil.

Before each fMRI session, monkeys received an intravenous injection of 8–10 mg/kg monocrystalline iron oxide nanoparticles (MION; Molday ION, BioPAL) through the femoral or saphenous vein to increase contrast-to-noise ratio and reduce contributions from superficial draining veins^54^. To limit excessive iron accumulation, deferoxamine mesylate (Desferal; 1 g/day, intramuscularly) and deferiprone (Ferriprox; 60 mg/kg/day, orally) were administered after scanning for 20 days, until serum iron and ferritin levels returned approximately to baseline ranges.

### fMRI visual stimulation and experimental design

Visual stimuli were projected onto a translucent screen using a Barco LCD projector operating at 60 Hz with a resolution of 1400 × 1050 pixels. The screen was positioned 57 cm from the monkey’s eyes. Eye movements were monitored using an infrared corneal-reflection eye-tracking system (ISCAN, 120 Hz).

All images (24 grayscale images per category) were matched to one another and to the background in luminance, gamma-corrected and controlled for spatial frequency (SHINE toolbox^56^). Images were presented centrally and were approximately 8.5° × 8.5°. The three inanimate object categories had been validated previously with image-based analyses and behavioral ratings^57^: the categories did not differ significantly in physical shape and tools and other manipulable objects were rated as similarly graspable, whereas tools were distinguished by a greater perceived capacity to extend the body’s acting space.

The experiment used a block design comprising seven stimulus conditions. Each stimulus block lasted 24 s. Condition order was counterbalanced across runs using a Latin-square procedure such that each condition occurred equally often at each serial position, yielding seven sequence orders. Each run began and ended with a fixation-only baseline block consisting of a uniform gray screen matched in luminance to the stimulus background. An additional fixation block was presented in the middle of the run to separate the two halves of the sequence, as illustrated in Fig. 1B.

Only runs with fixation performance exceeding 95% were included in the analysis. This resulted in 121 runs across the three monkeys: 46 runs for M1, 40 runs for monkey M2, and 35 runs for M3.

### fUSI Surgical preparation and acquisition

Two male rhesus macaques (M4, M5) participated in the fUSI experiments. Each animal was implanted with an MR-compatible headpost and a custom cranial recording chamber providing access to the parieto-occipital cortex, specifically, the cortical regions surrounding the superior temporal sulcus and intraparietal sulcus and was oriented approximately 24° from vertical in the coronal plane and 20° from vertical in the sagittal plane.

Chamber placement was verified using a computed tomography scan acquired with 0.4-mm slice thickness, 64 × 0.6-mm collimation, and a pitch of 0.85, with reconstruction using both bone and soft-tissue kernels. The CT volume was aligned to the average of ten high-resolution T1-weighted anatomical scans (0.4-mm isotropic resolution; TR = 2700 ms; TE = 2.5 ms; 3 T Siemens) using the Advanced Normalization tools registration module implemented in 3D Slicer^58,59^.

During recording, monkeys were seated in a primate chair with the head stabilized by the implanted headpost. Visual stimuli were presented on a screen positioned 40 cm from the eyes. Eye position was continuously monitored, and successful fixation was reinforced with liquid reward.

Functional ultrasound data were acquired using a volumetric fUSI probe connected to a Verasonics acquisition system. Volumes were acquired at 2 Hz, corresponding to one volume every 500 ms, with a spatial resolution of 100 × 150 × 150 µm³. The acquisition field of view covered approximately 1 cm² in-plane^26^.

### fUSI Visual stimuli and task design

The fUSI experiment used the same general class of static visual stimuli as the fMRI experiment, with the additional distinction between human and monkey body-part categories. Stimuli were approximately 8.5° in size.

The stimulus comprised nine categories: human hands, human faces, human bodies, monkey hands, monkey faces, monkey bodies, tools, manipulable objects, and non-manipulable objects. A fixation-only condition served as the control condition.

Each run began and ended with a 15-s fixation baseline. Between these baseline periods, five repetitions of the complete experimental sequence were presented. Each repetition contained ten blocks presented in pseudorandom order: nine stimulus-category blocks and one fixation-control block. Each block lasted 15 s. During stimulus blocks, visual images were presented for the first 4 s, followed by 11 s of fixation. During the 4-s stimulation period, eight images from the same category were presented sequentially for 500 ms each. Fixation-control blocks contained no image presentation.

Monkey M4 completed 95 runs across 13 recording sessions, corresponding to 475 complete repetitions, whereas monkey M5 completed 109 runs across 8 sessions, corresponding to 545 repetitions. The final fUSI dataset therefore comprised 204 runs and 1,020 complete experimental repetitions.

### Image Similarity Calculation

To quantify low-level image similarity among the stimulus categories, we computed the Structural Similarity Index (SSIM) on both the original images and their outlines using the structural_similarity function from skimage.metrics ^60^. SSIM provides a robust measure of structural integrity. Extended data figure 2 shows the resulting heatmaps for the original stimuli and their outlines. The results indicate that tools, manipulable objects, and non-manipulable objects exhibit high similarity.

## Data analysis

### fMRI image reconstruction and preprocessing

We used the same procedures as in our previous studies^24,61^, which will be briefly summarized. To decrease artefacts caused by monkey movements, the “optimized generalized autocalibration partially parallel acquisitions” (optimized GRAPPA) reconstruction method was used to reconstruct the EPI images. We used the Bioimage suite: http://bioimagesuite.yale.edu and Freesurfer http://surfer.nmr.mgh.harvard.edu for preprocessing. After skull stripping, motion correction with 6 degrees of freedom was performed on the EPIs, using one EPI volume from the anesthesia scan session as template. To reduce frame-to-frame distortions, a slice-by-slice undistortion correction procedure using a B-spline grid-based nonlinear registration method was performed, according to the same template.

General linear model (GLM) analysis

The GLM was fitted in Freesurfer to calculate t-score and percent signal change maps. Four volumes collected at the start of each run served as dummy scans and were discarded. Both linear and second-order regressors centered around zero were included in the model to remove low-frequency scanner signal drifts. To account for physiological and other nuisance temporal domain noise, a GLM denoising algorithm was performed based on white matter and ventricle signals to generate additional noise regressors. The condition regressors were generated by convolving the predictors by a canonical gamma fit monocrystalline iron oxide nanoparticle (MION) hemodynamic response function (HRF).

### Surface segmentation and reconstruction

Cortical surface reconstruction followed the non-human primate version of the Human Connectome Project (HCP) pipeline, combining FSL, FreeSurfer, and Connectome Workbench (https://humanconnectome.org). Because all reference anatomical images were acquired in a single anesthetized session with the head fixed by the headpost, movement between T1-weighted and T2-weighted volumes was negligible; we therefore omitted motion correction to avoid unnecessary smoothing. T2-weighted images served to exclude cerebrospinal fluid and pial veins during surface segmentation.

### Registration between EPI and structural images, and cortical surface projections

EPI distortions induced by magnetic-field inhomogeneities were corrected as in our previous studies^55^: a distortion field computed from the field maps of the same anesthesia session was applied to the GLM results with FSL. Using the T1-weighted volume acquired in that anesthetized session as intermediate space, functional results were registered to the high-resolution reference anatomical images. For surface projection we retained only voxels located between 20% and 80% of cortical depth (between pial surface and grey–white matter boundary), ensuring signals originate from grey matter and avoiding cross-talk between neighboring banks of the same sulcus; data were not smoothed.

### Region of interest definition

Functional regions of interest (fROIs) were defined on a localization dataset comprising 60 of the 121 runs (split 1), kept strictly independent from the remaining 61 runs (split 2) used for all response-profile and representational analyses. Functional data from all three monkeys were registered to the MEBRAINS template space^28^, which served as the common space for fROI definition and anatomical localization, with cortical labels assigned using a macaque atlas implemented in that space. Left-and right-hemisphere fROIs were not analyzed separately.

Several sets of fROIs were defined for different analyses. First, animate-and inanimate-preferring fROIs were identified using the Animate > Inanimate and Inanimate > Animate contrasts, where the animate category comprised face, body, and hand, and the inanimate category comprised tool, manipulable objects (Mani), non-manipulable objects, and chair. Second, category-preferring fROIs were defined separately for each of the seven stimulus categories using category-versus-all-other-categories contrasts (e.g., hand > face + body + tool + manipulable objects + non-manipulable objects + chair). These fROIs were thresholded at *p* <.0001, uncorrected.

To characterize hand-related organization more specifically, two additional contrasts were used: hand > face + body and hand > inanimate (tool + manipulable objects + non-manipulable objects + chair). The union of the two thresholded maps was used to define broader hand-related Union fROIs, whereas their spatial intersection was used to define more restrictive Intersection fROIs. These fROIs were subsequently used to evaluate category-response profiles using the independent dataset (split 2).

For the spatial-overlap analysis, pair-specific fROIs were defined separately for each animate–inanimate category pair. To avoid trivially favoring overlap between the two categories being compared, the paired category was excluded from the reference condition of each localizer. For example, hand–tool overlap was evaluated between a hand > others excluding tool fROI and a tool > others excluding hand fROI. Pair-specific fROIs were thresholded at *p* < 0.001, uncorrected. The detailed overlap procedure is described below.

Finally, hand-preferring fROIs used for the Crossnobis RDM and brain–model RSA analyses were defined using the hand > others excluding tool (*p* < 0.001) from the same localizer dataset (split 1). All subsequent multivoxel representational analyses within these fROIs were performed using split 2 data.

### Activation profiles and statistical comparisons

Category-response profiles within the hand-related fROIs were evaluated on the 61 runs (split 2) held out from functional localization. Analyses were performed separately for the Union and Intersection fROIs defined from the hand > face + body and hand > Inanimate localizers. For each run and each of the seven stimulus categories (face, body, hand, tool, manipulable objects, non-manipulable objects, and chairs), the condition-specific response estimate relative to fixation was averaged across all voxels within the corresponding fROI, yielding one response value per condition and run. fROIs containing 10 or fewer voxels were excluded from these response-profile analyses.

For visualization, condition responses were averaged across runs, and error bars indicate the standard error of the mean (SEM) across runs. Statistical differences among category responses were assessed using paired two-tailed t-tests across runs for all 21 pairwise comparisons among the seven conditions. The resulting p values were corrected separately within each fROI across the 21 pairwise tests using the Benjamini–Hochberg false discovery rate (FDR)^62^. FDR-adjusted p values below.05 were considered significant. The hand-centered statistical summaries reported in the figures were taken from the six comparisons between hand and each of the remaining stimulus categories within this same set of FDR-corrected pairwise tests.

Finite impulse response (FIR) time courses were additionally examined in the same independently defined Union and Intersection fROIs using split 2 data. Condition-specific β estimates were summarized at each time point to visualize the temporal response profile of the seven stimulus categories within each fROI.

### Spatial overlap analysis

To determine whether individual inanimate categories occupied spatially overlapping functional territories with hand-, face-, or body-related representations, pairwise overlap analyses were performed between each of the three animate categories (hands, faces, and bodies) and each of the four inanimate categories (tools, manipulable objects, non-manipulable objects, and chairs).

To avoid circularly favoring overlap between the two categories being compared, fROIs were defined separately for each category pair using pair-specific contrasts in which the paired category was excluded from the reference set. For example, hand–tool overlap was evaluated between a hand fROI defined by hand > other categories excluding tool and a tool fROI defined by tool > other categories excluding hand. The same procedure was applied independently to all animate–inanimate category pairs. Consequently, the hand, face, or body fROI used in one pairwise comparison was not necessarily identical to that used in another comparison. Pair-specific fROIs were thresholded at p <.001 (uncorrected) for this analysis.

Overlap was evaluated within atlas-defined cortical areas. Category pairs were included only when both corresponding pair-specific fROIs contained more than 15 voxels. For each eligible pair, spatial overlap was first quantified as the absolute number of voxels belonging to both fROIs:

Overlap size = |A ∩ B|

where A and B denote the two pair-specific fROIs.

Because the absolute number of overlapping voxels is influenced by the sizes of the two fROIs, proportional overlap was additionally quantified using the overlap coefficient:

Overlap coefficient = |A ∩ B| / min(|A|, |B|)

The overlap coefficient expresses the proportion of the smaller fROI that is shared with the larger fROI. A value of 0 indicates no spatial overlap, whereas a value of 1 indicates that the smaller fROI is completely contained within the larger fROI. Both absolute overlap size and the overlap coefficient were used to characterize the spatial relationship between animate and inanimate category representations.

### Crossnobis representational dissimilarity analysis

To examine category relationships at the multivoxel representational level, representational dissimilarity matrices (RDMs) were calculated within independently defined hand-preferring fROIs (split 1). The remaining split 2 data were used for all Crossnobis analyses. fROIs containing 10 or fewer voxels were excluded.

For each fROI and run, voxel-wise response estimates relative to fixation were extracted separately for the seven stimulus categories, yielding a condition-by-voxel activation pattern. Representational dissimilarity between category pairs was quantified using a cross-validated Mahalanobis distance (the Crossnobis)^63,64^ with leave-one-run-out cross-validation. Crossnobis distance has also been applied to multivoxel representational analyses in non-human primate fMRI^65^.

Noise precision was estimated separately within each fROI. For each pair of categories and each held-out run, the difference between their multivoxel response patterns in the held-out run was cross-validated against the corresponding difference between their mean response patterns across all remaining runs. A diagonal noise-precision matrix was estimated within each fROI from residual variability pooled across runs and conditions. The run-wise Crossnobis distance was calculated as:

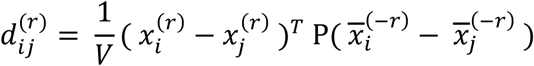

Where *x_i_*^(*r*)^ denotes the multivoxel response pattern for category in the held-out run r, 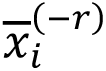 denotes the mean response pattern for category across all runs except run r, P is the diagonal noise-precision matrix, and V is the number of valid voxels contributing to the comparison.

This procedure yielded one symmetric 7 × 7 Crossnobis RDM for each run and fROI. The RDM shown for each fROI was obtained by averaging the corresponding run-wise Crossnobis distances across the split 2 data. Because Crossnobis distances are cross-validated, their estimates are not constrained to be positive. Values around zero indicate little reproducible differentiation between two category patterns, whereas positive values indicate increasing representational dissimilarity.

For the pairwise distance summary, each of the 21 unique run-wise Crossnobis distances was additionally tested against zero using a one-sample sign-flip permutation test across runs^66^. Multiple comparisons were controlled separately within each fROI across the 21 category pairs using the Benjamini– Hochberg FDR procedure.

### Pairwise Crossnobis distance comparisons

To characterize the hand–tool representational relationship, we examined six Crossnobis distances within each independently localized hand-preferring fROI: face–tool, body–tool, hand–tool, hand–manipulable objects, hand–non-manipulable objects, and hand–chair. The hand–tool distance was compared with each of the other five distances using paired-sample tests across the runs of split 2. The resulting P values were corrected for the five comparisons within each fROI using the Benjamini–Hochberg false discovery rate (FDR) procedure. For the summary across all hand-preferring fROIs, the 12 distances relating hands, bodies, and faces to the four inanimate categories were tested against zero using one-sided sign-flip permutation tests with 10,000 permutations. Within each fROI, P values were FDR-corrected across the 12 category-pair comparisons. FDR-adjusted P values below 0.05 were considered significant.

### Computational model representations

To examine whether different types of learned visual representations captured the category geometry observed in the neural RDMs, four candidate predictor RDMs were used: CORnet-V1, CORnet-IT, and representations derived from Video Swin Transformer models trained on Kinetics-400 (K400) and Something-Something V2 (SSV2). All model RDMs contained the same seven categories and category ordering as the neural RDMs.

### CORnet-S

CORnet-S was used to characterize representations associated with static object recognition^35,36^. Stimulus images were resized to 224 × 224 pixels and normalized using ImageNet normalization parameters before being presented individually to the pretrained network.

Representations were extracted from the V1 and IT stages of CORnet-S, providing relatively early and higher-level visual representations, respectively. For each stimulus image, feature maps were flattened across channels and spatial dimensions (C × H × W) into a single feature vector, retaining activations at each spatial location. Feature vectors were then averaged element-wise across stimulus exemplars belonging to the same experimental category, yielding one category-level representation for each of the seven categories. Pairwise dissimilarities between category-level representations were quantified using correlation distance, defined as 1 − Pearson correlation, yielding separate 7 × 7 RDMs for the V1 and IT stages, hereafter referred to as CORnet-V1 and CORnet-IT.

### Action-trained video models

To characterize representations learned from action-recognition data, pretrained Video Swin Transformer models^39^ trained on Kinetics-400^37^ and Something-Something V2^38^ were used. Kinetics-400 contains a broad range of human action categories, whereas Something-Something V2 emphasizes fine-grained, temporally structured human hand–object interactions.

Because the experimental stimuli were static images, each image was repeated across 32 identical frames to form a static video clip before being presented to the video models. No additional motion information was therefore introduced at test time. Feature activations from the Video Swin backbone were averaged across temporal and spatial dimensions to obtain one feature vector for each stimulus. Feature vectors were subsequently averaged across exemplars within each of the seven experimental categories, and pairwise correlation distances were used to construct separate 7 × 7 K400 and SSV2 RDMs.

### Brain–model representational similarity analysis

The correspondence between each candidate model and the neural representational geometry was first quantified using representational similarity analysis (RSA^5^). For each fROI and run, the 21 unique pairwise dissimilarities from the lower triangle of the 7 × 7 neural Crossnobis RDM were vectorized in the same category-pair order as the corresponding model RDM. Spearman rank correlation was then calculated between the neural and model RDM vectors, yielding one RSA coefficient (Spearman’s ρ) for each fROI, run, and model.

Run-wise Spearman correlation coefficients were averaged within six approximately equal, non-overlapping groups of consecutive runs. Positive brain–model correspondence was assessed using a one-sided exact sign-flip test of the group-level estimates against zero. Reported RSA coefficients represent the mean of these group-level estimates. Multiple comparisons were controlled using the Benjamini–Hochberg FDR procedure across fROI–layer combinations separately within each model, with FDR-adjusted p-values below.05 considered significant.

Because the four candidate predictor RDMs shared some representational structure, a multiple-regression RSA was additionally performed to estimate the contribution of each predictor while accounting for the others. CORnet-V1, CORnet-IT, K400, and SSV2 were entered simultaneously into an ordinary least-squares regression model. For each fROI and run, the 21 neural Crossnobis distances formed the dependent variable and the corresponding 21 dissimilarities from the four predictor RDMs formed the independent variables. Before regression, the neural RDM vector and each predictor RDM were independently standardized across the 21 category pairs to zero mean and unit variance. The regression therefore yielded one standardized coefficient (β) for each predictor, run, and fROI. Because all four predictors were included simultaneously, each standardized β quantified the contribution of that predictor to the neural representational geometry after accounting for representational variance shared with the remaining predictors.

For visualization, standardized β coefficients were averaged across runs, with error bars representing the SEM across runs. Evidence for a positive independent contribution was assessed for each predictor using a one-sided sign-flip permutation test of the run-wise β coefficients against zero with 10,000 permutations. The resulting p values were corrected for multiple comparisons using the Benjamini–Hochberg FDR procedure.

### VIFs: Variance Inflation factors

Multicollinearity among the four model RDM predictors was assessed using variance inflation factors (VIFs). The 21 unique off-diagonal dissimilarities from each model RDM were standardized across condition pairs. For each predictor, an auxiliary ordinary least-squares regression with an intercepted predicted value from the remaining three predictors. VIF was calculated as 1/(1 − R²), where R² denotes the coefficient of determination of the auxiliary regression. VIF values of 5 and 10 were used as descriptive reference levels.

The full four-predictor model was retained as the regression analysis. To assess sensitivity to predictor inclusion, regression models were refitted separately for each fROI and run after omitting one predictor at a time. All model sets used identical fROIs, runs, and valid condition pairs, with the same standardization parameters for shared predictors and neural RDMs. Sensitivity was characterized by changes in coefficient sign and magnitude and in the pattern of mean standardized β coefficients across fROIs. Particular attention was given to SSV2 coefficients after separately omitting CORnet-V1, CORnet-IT, or K400. Coefficients were visualized as means ± SEM across matched runs of split 2 per fROI.

### First-level functional Ultrasound (fUSI) Data Analysis

First-level statistical analyses were performed using the General Linear Model (GLM) implemented in the Nilearn Python library. To accurately capture the specific vascular dynamics of functional ultrasound imaging, task regressors were convolved with a custom fUSI-specific Hemodynamic Response Function (HRF)^26^. This custom HRF was peak-normalized and interpolated to match the acquisition repetition time (TR = 0.5 s).

The GLM was fitted using an Ordinary Least Squares (OLS) noise model without a cosine drift, and spatial smoothing was applied using a Gaussian kernel with a Full-Width at Half-Maximum (FWHM) of 0.2 mm. To control for gaze-related artifacts, eye-tracking coordinates (X and Y gaze positions) were included in the design matrix as nuisance regressors.

The explicit baseline (’fixation’) was kept in the model. We computed three sets of statistical contrasts to evaluate the evoked hemodynamic responses: (1) single-condition contrasts against the fixation baseline, (2) pairwise comparisons between all individual stimulus conditions. For each contrast, statistical parametric maps, effect sizes (beta estimates), and effect variances were systematically generated.

### Categorical Selectivity and Winner-Take-All (Gradation) Analysis

To identify voxels displaying high functional selectivity for specific categories, a voxel-wise Winner-Take-All (WTA) or gradation analysis was implemented on the first-level general linear model (GLM). For each of the nine experimental conditions (human bodies, monkey bodies, human faces, monkey faces, human hands, monkey hands, manipulable objects, non-manipulable objects, and tools), a target condition was systematically compared against the remaining eight conditions through individual pairwise fixed-effects contrasts. A conservative statistical threshold of Z > 3.09 (equivalent to p < 0.001, uncorrected) was applied to each contrast. For each voxel, a cumulative gradation score ranging from 0 to 8 was computed by summing the binary hit masks across all eight pairwise comparisons. Voxels achieving a maximum score of 8 were classified as strict Winner-Take-All voxels, representing cortical sites where the target condition significantly outperformed every other experimental category.

## Data Availability

The data will be made available after acceptance of the manuscript.

## Code Availability

The custom codes will be made available after acceptance of the manuscript.

## Acknowledgments

The authors thank C. Fransen, I. Puttemans, A. Hermans, C Ulens, S. Verstraeten, W. Depuydt, and M. Depaep for technical and administrative support. This work received funding from KU Leuven C14/21/111 and C16/26/013, CELSA/24/020; the French national research agency under the reference ANR-20-CE37-0005; the European Union’s Horizon 2020 Framework Programme for Research and Innovation under Grant Agreement No 945539 (Human Brain Project SGA3); Fonds voor Wetenschappelijk onderzoek, Vlaanderen (12AF124N), and Grant Agreement No 856495 (European Research Council (ERC)).

## Contributions

Conceptualization: SB, HO, RF, QZ, WV. Methodology: YL, QZ, MF, CW, NF, GM, AU, DC, HW, XL, WV. Investigation: YL, MF, XL, QZ, WV. Funding acquisition: QZ, RV, WV. Supervision: QZ and WV. Writing – original draft: YL, QZ, WV. Writing – review & editing: YL, MF, CW, NF, GM, AU, DC, HW, XL, HO, SB, RV, QZ, WV.

## Competing Interests Statement

Authors declare that they have no competing interests.

## Extended Data Figures

**Extended data Figure 1.**
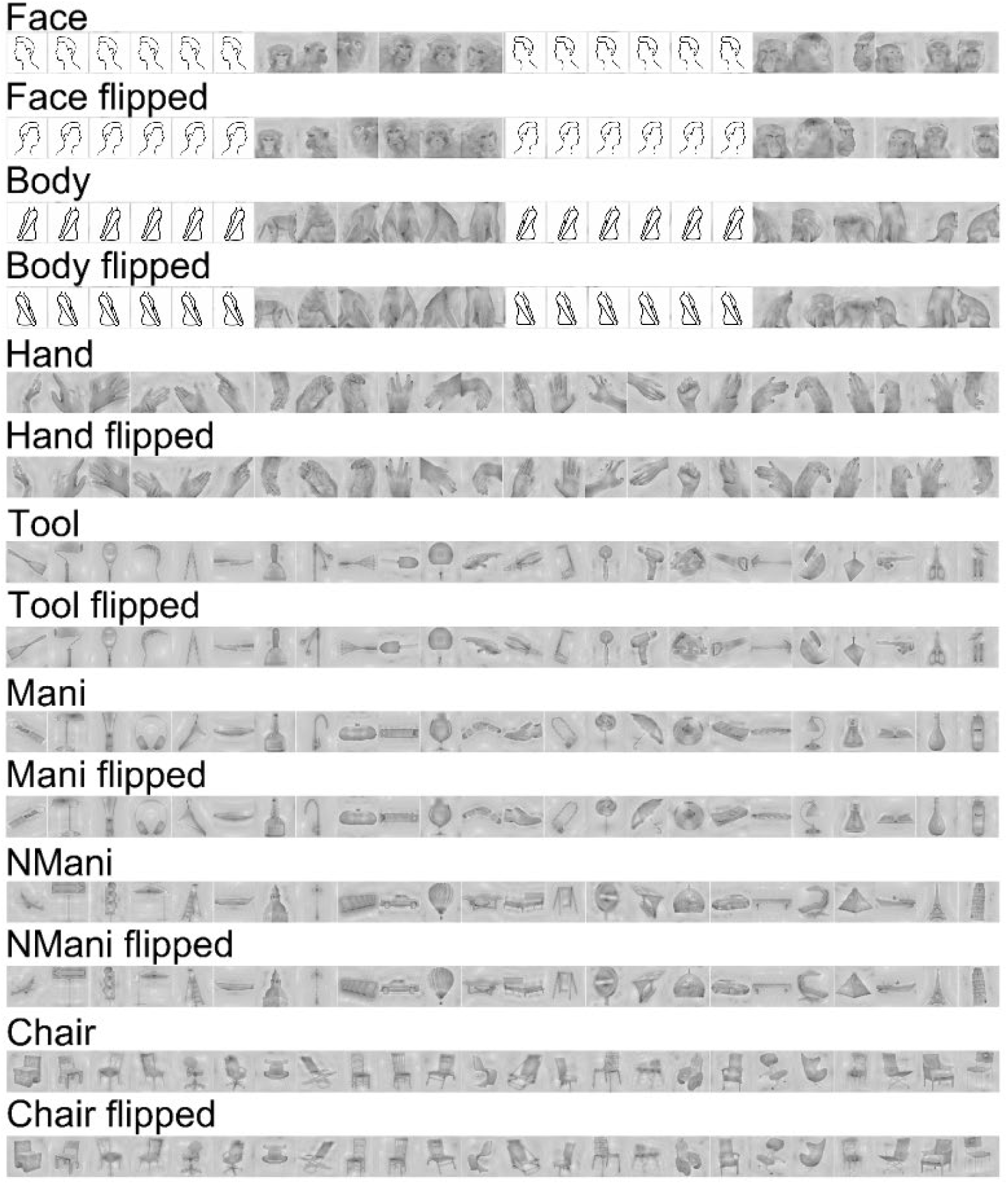
Stimulus set. The stimulus set was similar to, but not identical to, that used by^10,11^. The stimuli comprised seven categories: faces, bodies, hands, tools, manipulable objects (Mani), nonmanipulable objects (NMani), and chairs. Each category consisted of 24 original images and their horizontally flipped versions (48 images per category). Human faces and bodies are depicted as icons. The original images of humans used in the experiments are available upon request.

**Extended data Figure 2.**
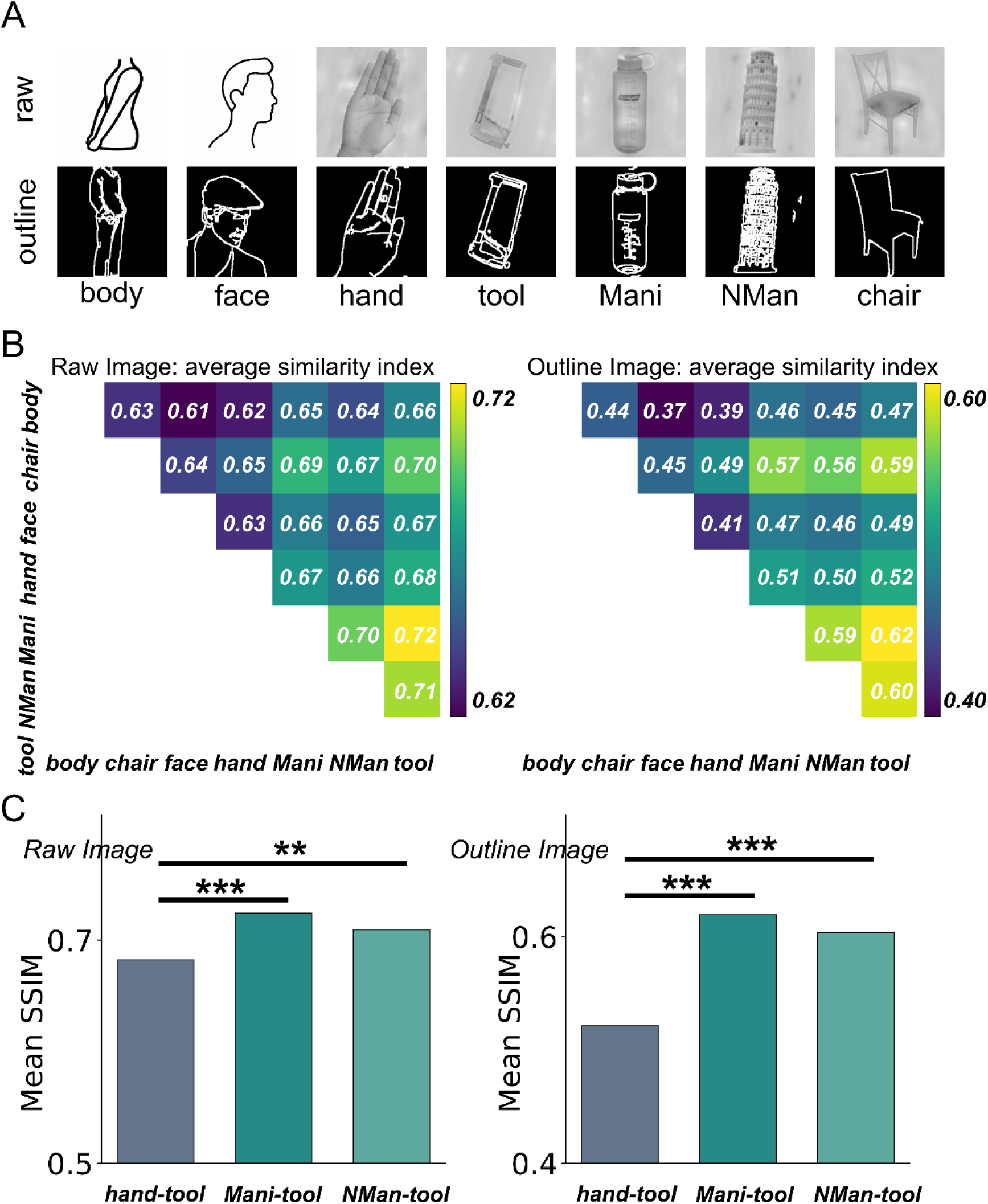
Stimulus similarity. A. Low-level similarity of the stimulus categories. Examples of the original stimuli are shown on the left and the corresponding outlines on the right. Human faces and bodies are depicted as icons. The original images of humans used in the experiments are available upon request. B. Structural Similarity Index (SSIM), computed for the entire set of original images and for their corresponding outlines. Note that tools are more similar to manipulable objects, non-manipulable objects and chairs compared to hands (see methods). C. Comparison of mean Structural Similarity Index (SSIM) values between tools and hands, Mani, and NMan, calculated separately for the original grayscale images (left) and their corresponding outlines (right). Bars represent mean SSIM values. Statistical significance was assessed using paired permutation tests, treating each original tool image and its horizontally flipped version as a single stimulus identity (24 identities). P-values were corrected for multiple comparisons using the Benjamini–Hochberg false discovery rate (FDR) procedure. **p < 0.01, ***p < 0.001. Tools showed significantly greater structural similarity to Mani and NMan than to hands in both image formats.

**Extended data Figure 3.**
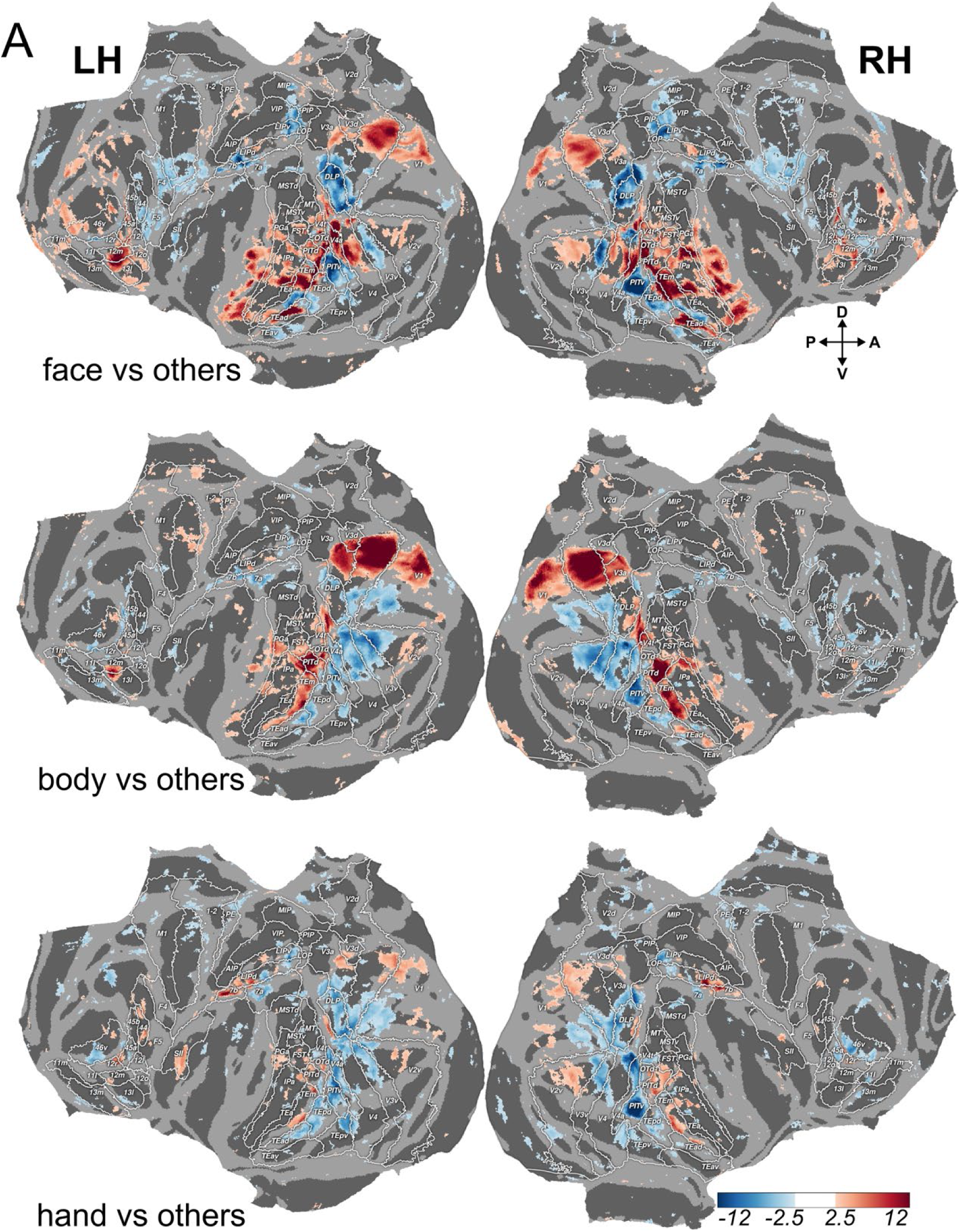
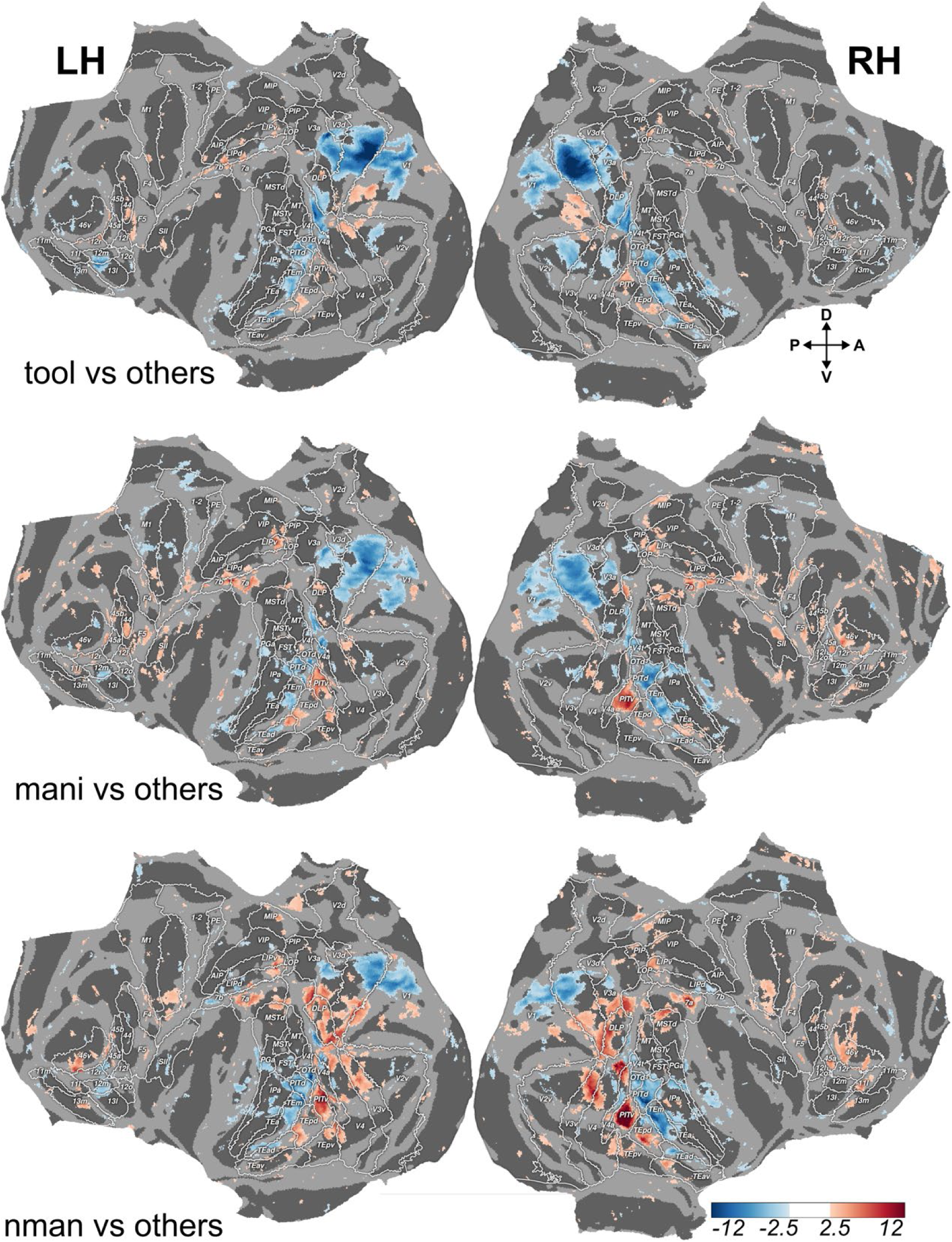
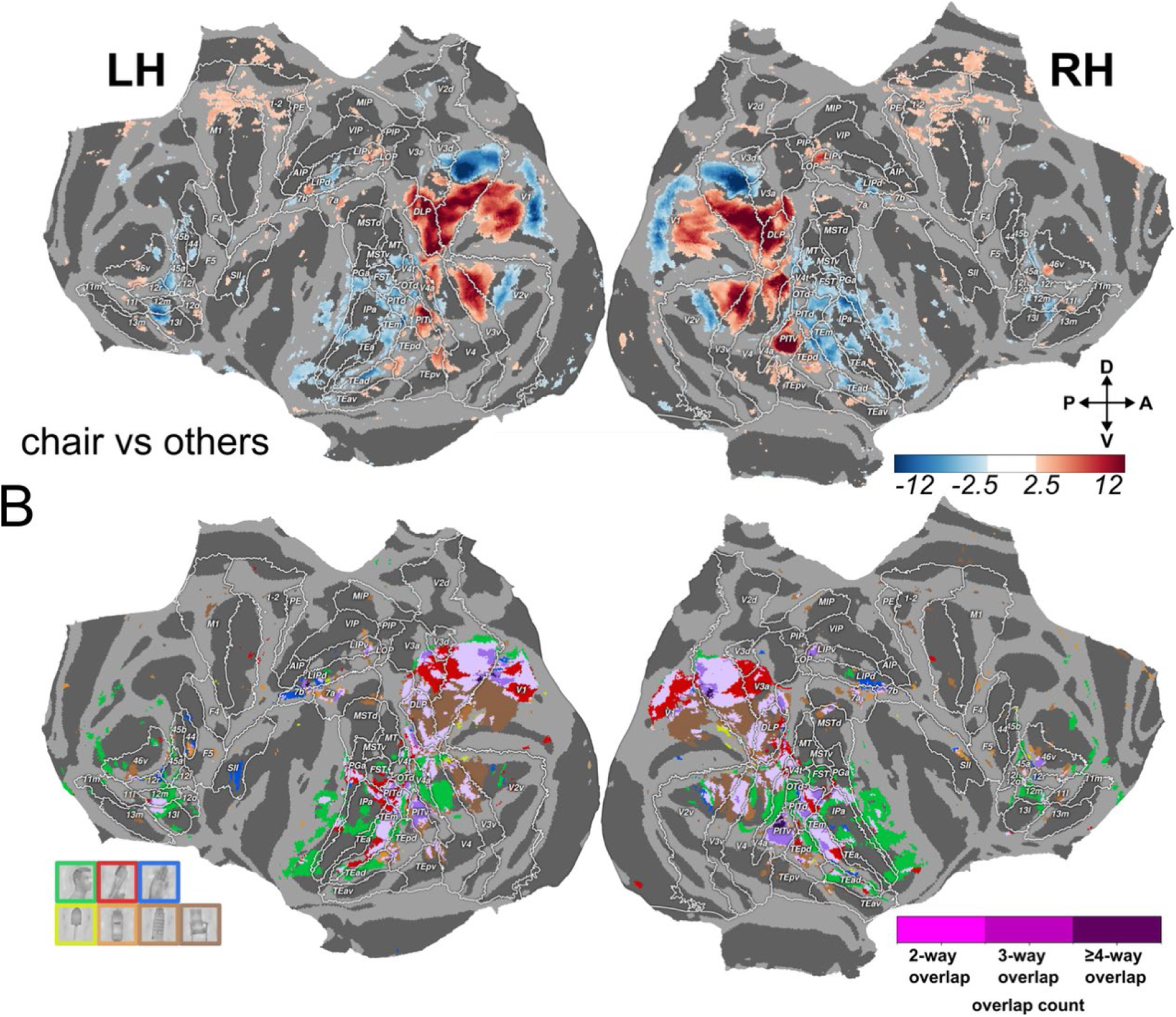
Whole-brain spatial distribution of category-preferences. A. Cortical surface maps for the seven category-versus-others contrasts: faces, bodies, hands, tools, manipulable objects, non-manipulable objects, and chairs. For each contrast, responses to the target category were compared with the average response across the other six categories (which we refer to as preference maps). T-score maps (threshold |t| > 2.5) are projected flattened views of the left and right hemisphere. B. Category-preferring fROIs were defined from split 1 using category-versus-others contrasts for faces, bodies, hands, tools, manipulable objects, non-manipulable objects, and chairs (p < 0.0001, |t| > 3.9). Colors indicate fROIs corresponding to each category. Pink indicates voxels shared across categories, with light, medium, and dark pink indicating voxels shared by two, three, or more categories, respectively.

**Extended data Figure 4.**
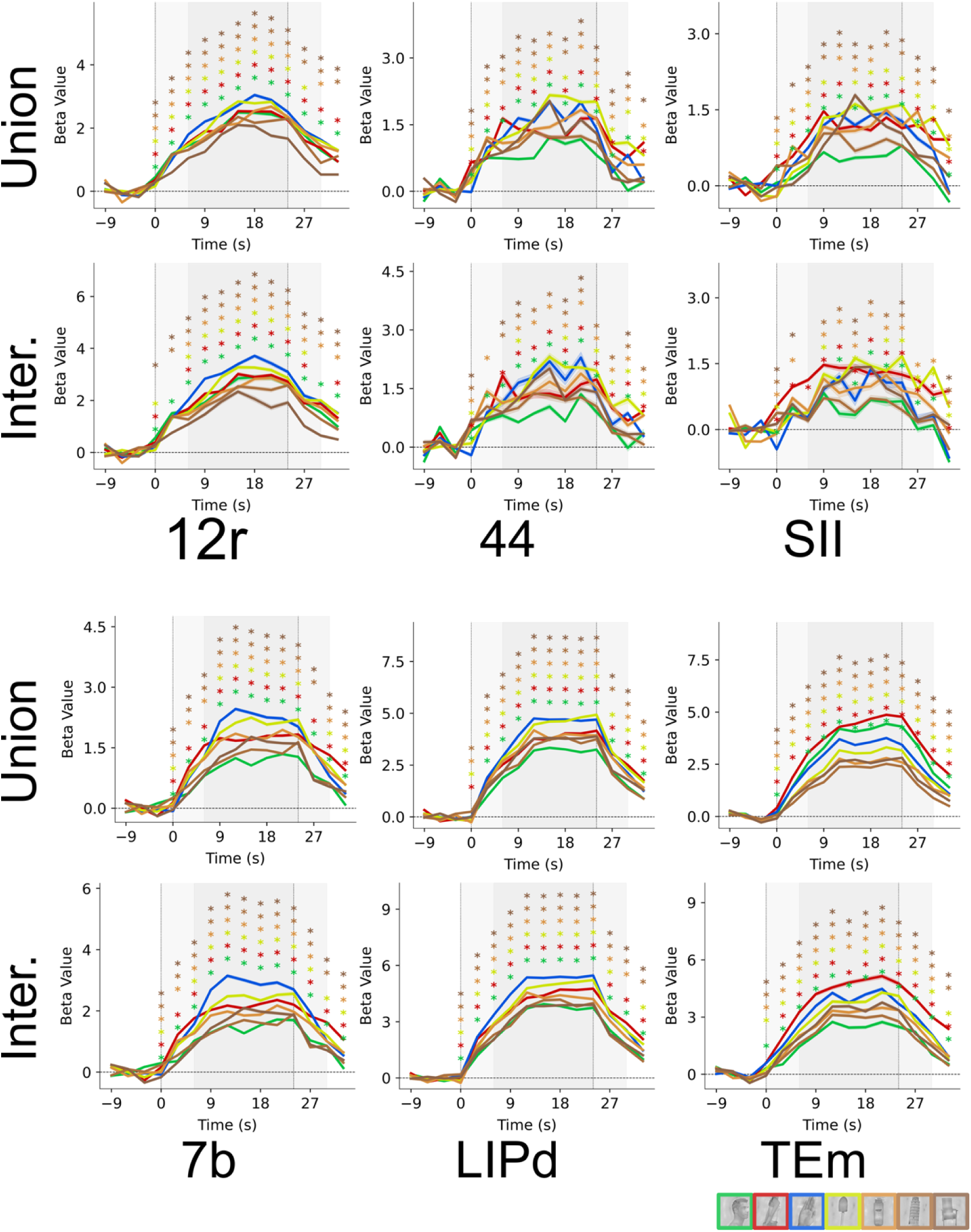
Event-related response time courses. Finite impulse response (FIR) time courses were estimated from the 61 runs (split 2) held out from fROI definition (split 1 data). Curves show mean FIR β estimates for each of the seven stimulus categories in areas 7b, LIPd and TEm, separately for the union (top) and intersection (bottom) fROIs. Time is aligned to stimulus block onset (0 s), and vertical lines mark block onset and offset. At each time bin (3s), the response to hands was compared separately with the response to each of the six other stimulus categories using paired t-tests. Asterisks indicate significant differences (FDR-adjusted P < 0.05).

**Extended data Figure 5.**
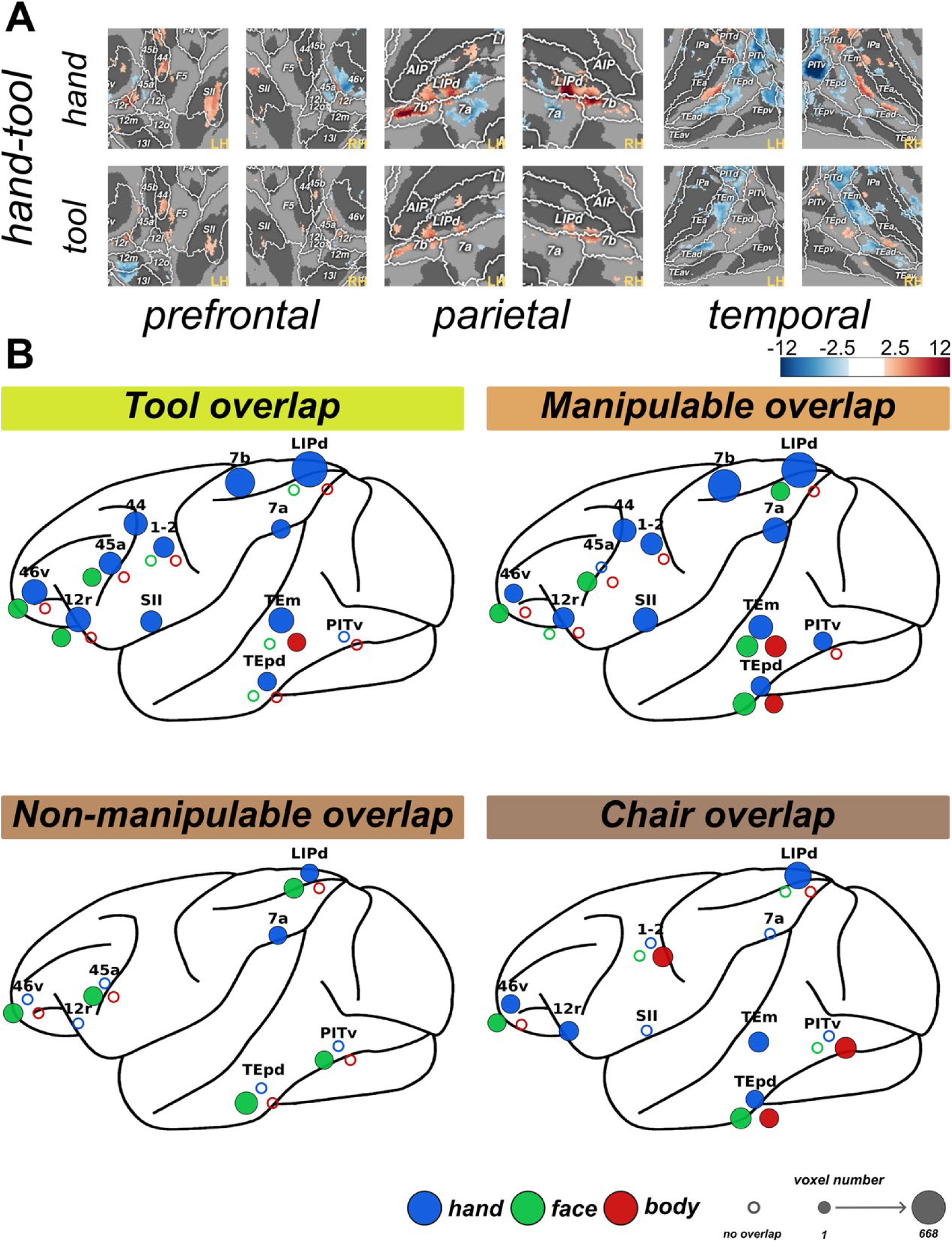
Spatial overlap between hand and tool representations. A. Representative sectors of flattened cortical t-score maps illustrating the pair-specific functional localizers used to assess hand–tool overlap in bilateral prefrontal, parietal, and inferotemporal cortex. Hand-and tool-responses were defined using the hand > others excluding tool, and tool > others excluding hand contrasts, respectively. White boundaries and labels indicate atlas-defined cortical areas. Warm and cool colors indicate positive and negative t values, respectively. For visualization, maps are displayed at (|t| > 2.5). B. Schematic of spatial overlap of tool, manipulable-object, non-manipulable-object, and chair localizers with hand-, body-, and face-sensitive regions. For each category pair, spatial overlap was defined as the intersection between the corresponding functional localizers (half data), with the paired category excluded from the control condition during fROI definition. For example, hand– tool overlap was defined as (hand > others excluding tool) ∩ (tool > others excluding hand). Functional localizers used for the overlap analysis were thresholded at (p < 0.001). Blue, red, and green circles indicate spatial overlap with hand-, body-, and face-sensitive ROIs, respectively. Circle size is proportional to the number of overlapping voxels (the quantitative data are presented in Extended data Table 2). Open circles indicate that both category-sensitive ROIs were present in the corresponding cortical region (anatomically-defined) but showed no spatial overlap. LH, left hemisphere; RH, right hemisphere.

**Extended data Figure 6.**
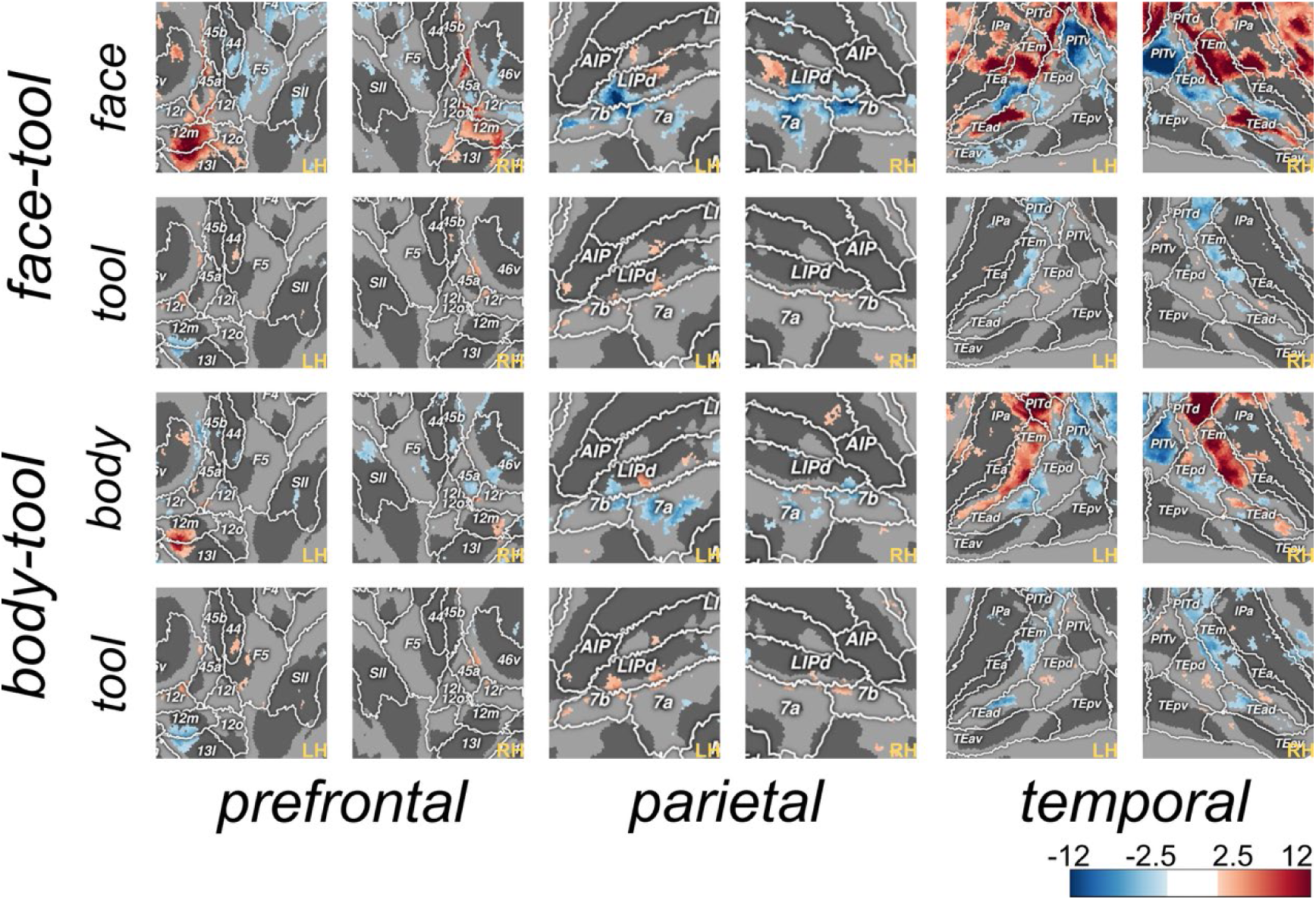
Spatial overlap between face/body and tool representations. Representative sectors of flattened cortical t-score maps illustrating the pair-specific functional localizers used to assess face/body–tool overlap in bilateral prefrontal, parietal, and temporal cortex. Face/body and tool-responses were defined using the face/body > others excluding tool, and tool > others excluding face/body contrasts, respectively. White boundaries and labels indicate atlas-defined cortical areas. Warm and cool colors indicate positive and negative t values, respectively. For visualization, maps are displayed at (|t| > 2.5).

**Extended data Figure 7.**
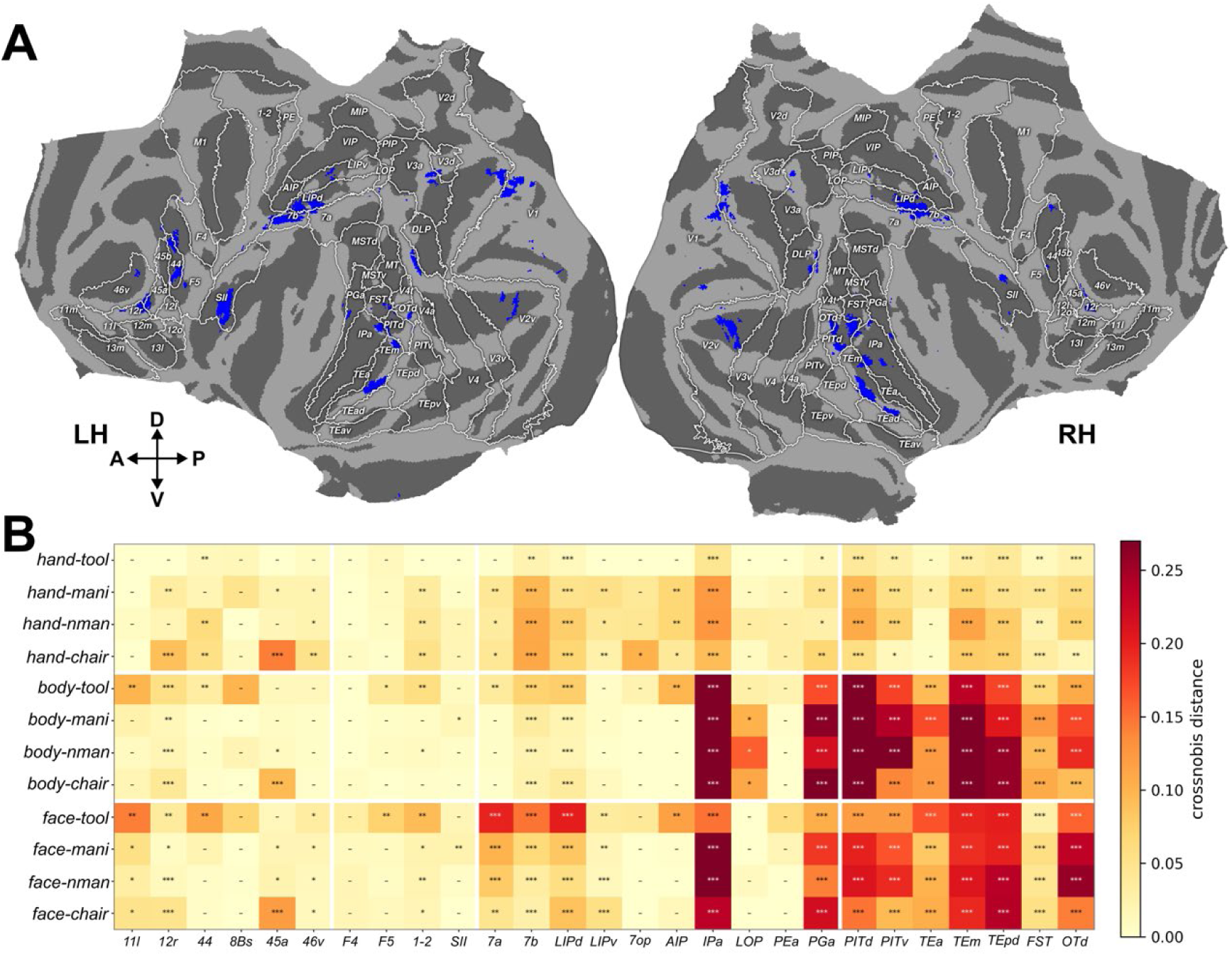
Pairwise representational distances across independently localized hand-preferring fROIs. A. Definition and cortical distribution of hand-preferring fROIs. Binarized t-score maps are shown on the flattened cortical surface for the left and right hemispheres. The fROIs were defined on the independent localizer dataset (split 1) using the contrast hand > mean of all other categories except tools at P < 0.001. B. Selected pairwise Crossnobis distances across hand-preferring fROIs. The heatmap shows Crossnobis distances between hands, bodies, and faces and each of the four inanimate categories (tools, manipulable objects, non-manipulable objects, and chairs). The fROIs are grouped into prefrontal, sensorimotor, parietal, and temporal sectors. Colors indicate the mean cross-validated Crossnobis distance across runs, with darker colors denoting greater representational dissimilarity. Each distance was tested against zero using a one-sided permutation test across runs. P values were corrected across the displayed category-pair comparisons within each fROI using the Benjamini-Hochberg false discovery rate (FDR) procedure. *, **, and *** indicate FDR-adjusted P < 0.05, P < 0.01, and P < 0.001, respectively; dashes indicate nonsignificant distances.

**Extended data Figure 8.**
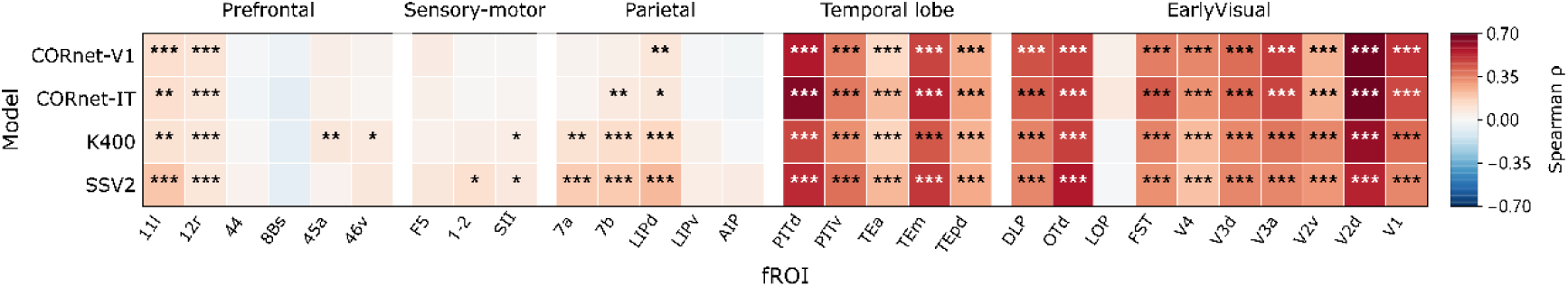
Model–brain RSA across hand-preferring fROIs. Spearman correlations between model RDMs and neural RDMs of hand-preferring fROIs. Neural RDMs were computed from the runs of split 2. Colors indicate mean Spearman correlation coefficients (ρ) across runs. Positive brain–model correspondence was assessed using one-sided sign-flip permutation tests of the run-wise correlations against zero (10,000 permutations). *, **, and *** indicate Benjamini–Hochberg FDR-adjusted P < 0.05, P < 0.01, and P < 0.001

**Extended data Figure 9.**
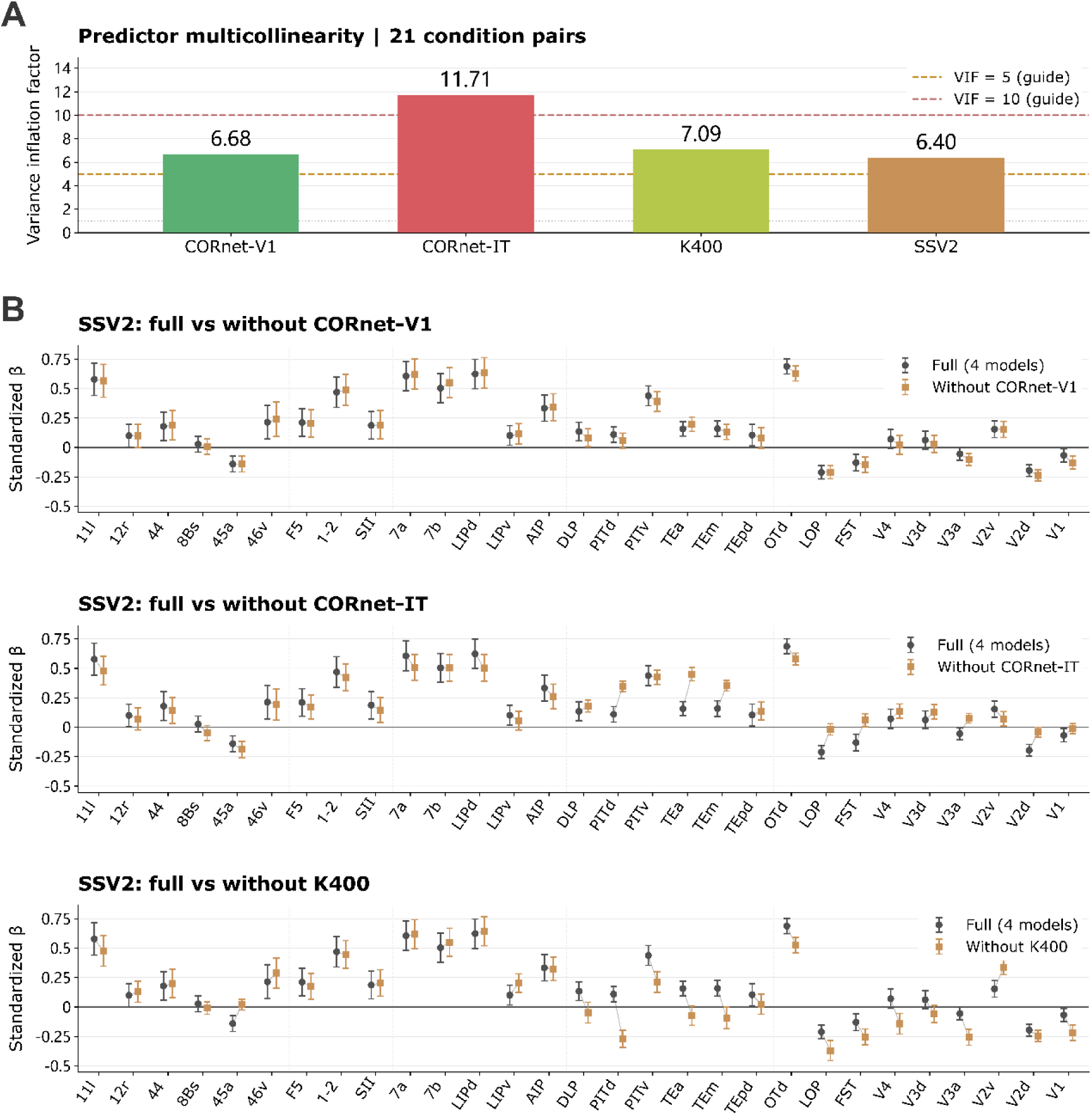
Multicollinearity diagnostics and sensitivity of SSV2 regression coefficients to predictor inclusion. A. Variance inflation factors (VIFs) for the four model RDM predictors, calculated from their 21 unique condition-pair dissimilarities. Dashed lines indicate VIF values of 5 and 10 as descriptive reference levels. B. Standardized SSV2 regression coefficients across fROIs from the full four-predictor model (gray circles) and reduced models (orange squares) omitting CORnet-V1, CORnet-IT, or K400, from top to bottom. Points and error bars indicate means ± SEM across 61 matched runs (split 2) per fROI. Thin gray lines connect the corresponding full-and reduced-model means. All comparisons used identical fROIs, runs, condition pairs, and standardization parameters.

**Extended data Figure 10.**
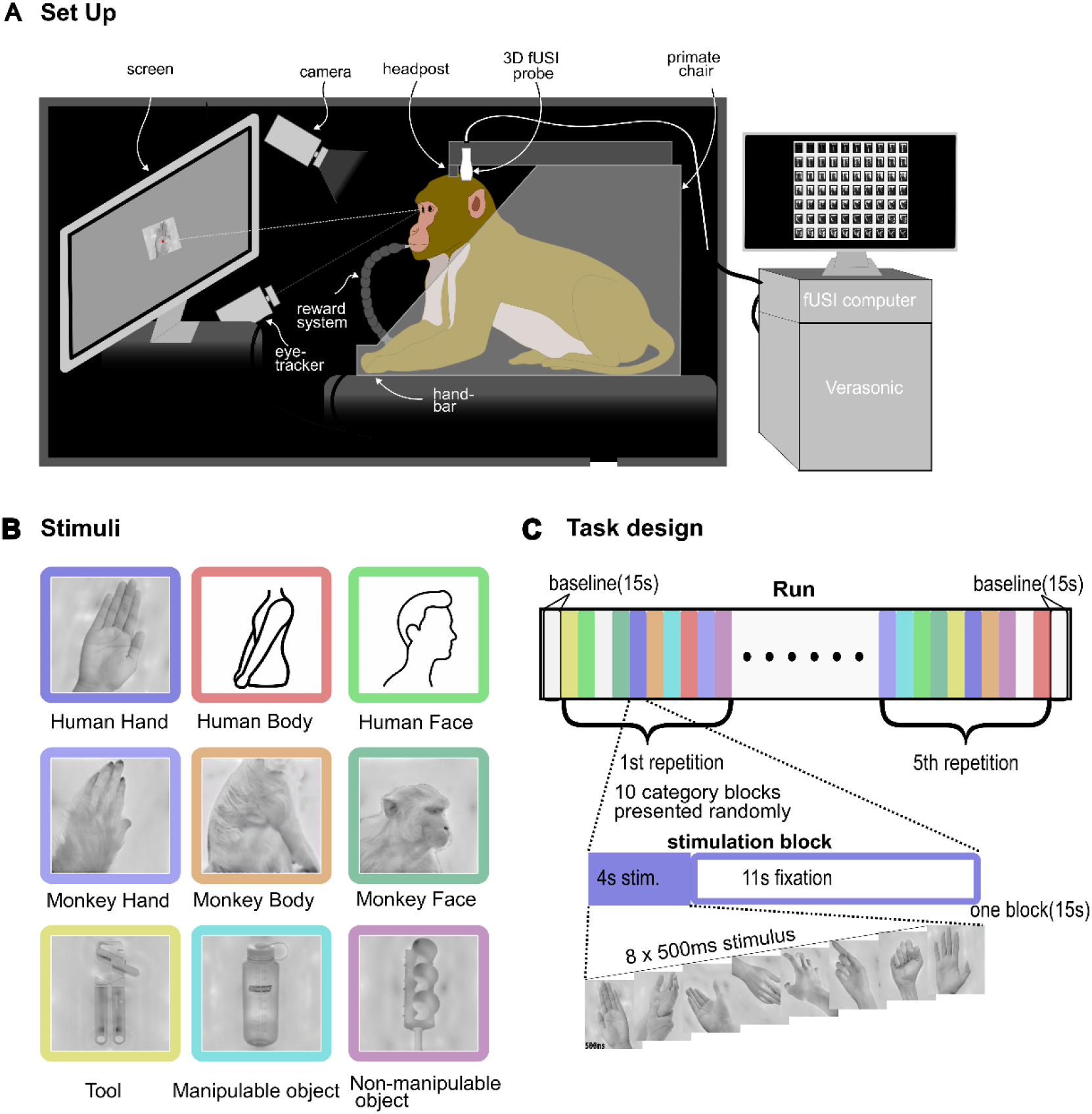
Experimental setup, visual stimuli, and task. design of the hand localizer for the 3D-fUSI experiments. A. Experimental setup for volumetric functional ultrasound imaging (3D-fUSI) in awake macaques. B. Stimulus categories used for the hand-localizer task: human hands, human bodies, human faces, monkey hands, monkey bodies, monkey faces, tools, manipulable objects, and non-manipulable objects. One representative example image is shown for each category. Human faces and bodies are depicted as icons. The original images of humans used in the experiments are available upon request. C. Block design of a single run. Each run began and ended with a 15-s fixation baseline. The complete set of stimulus categories was presented five times per run, with category order pseudorandomized within each repetition. Each category block lasted 15 s and comprised 4 s of visual stimulation followed by 11 s of fixation. During the stimulation period, eight images from the same category were presented sequentially for 500 ms each.

## Supplementary tables

**Extended data Table 1.**
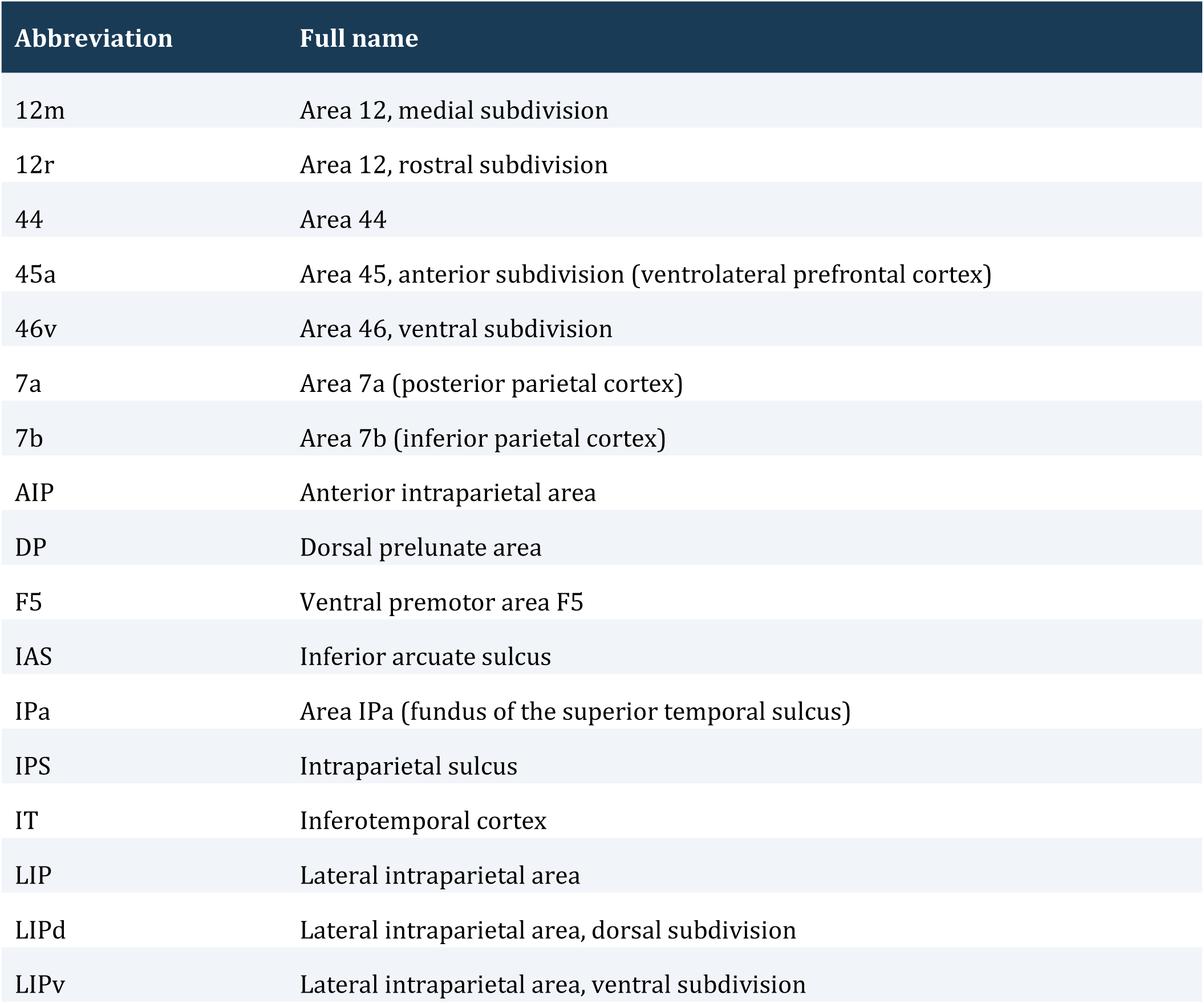
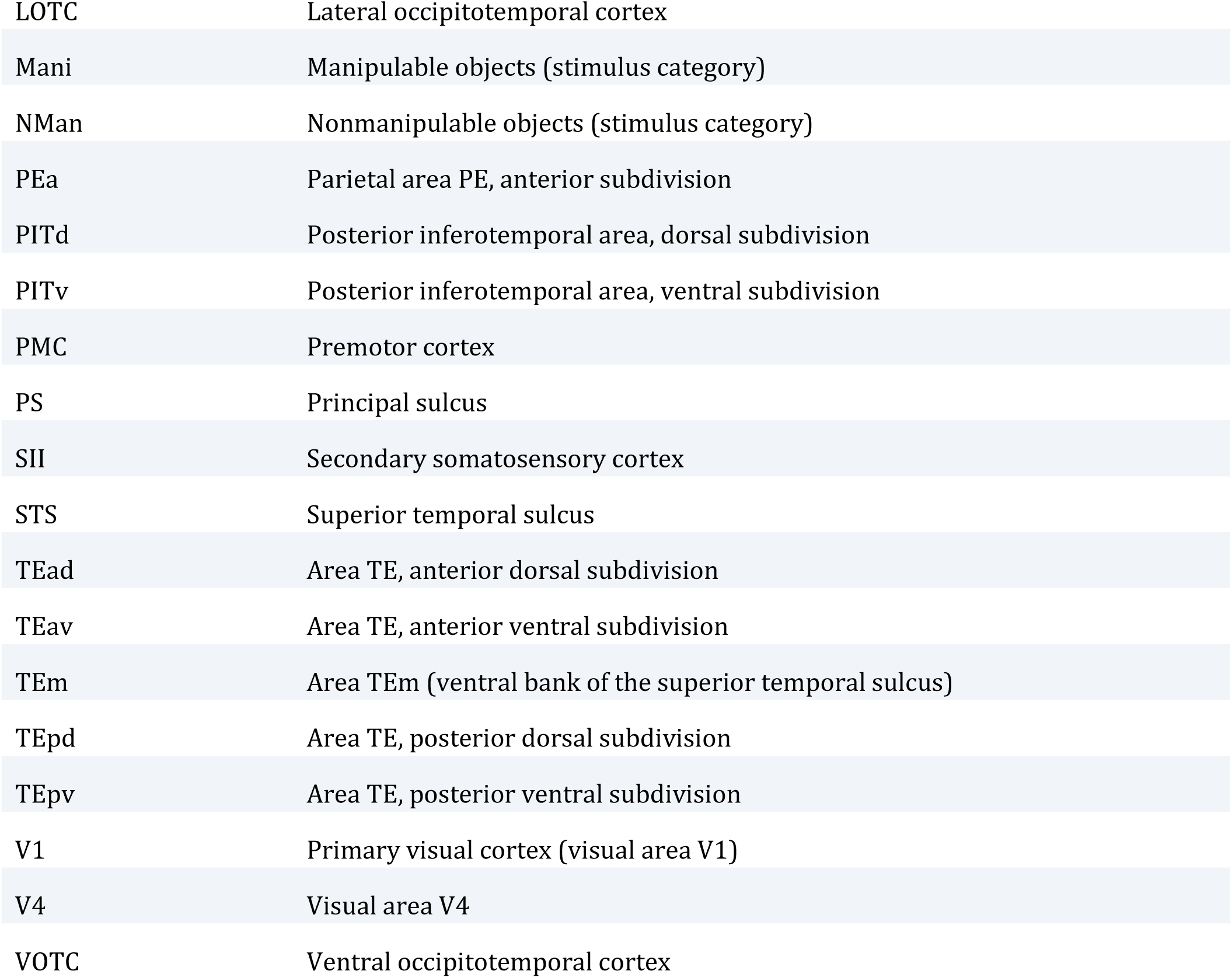
Abbreviations.

**Extended data Table 2.**
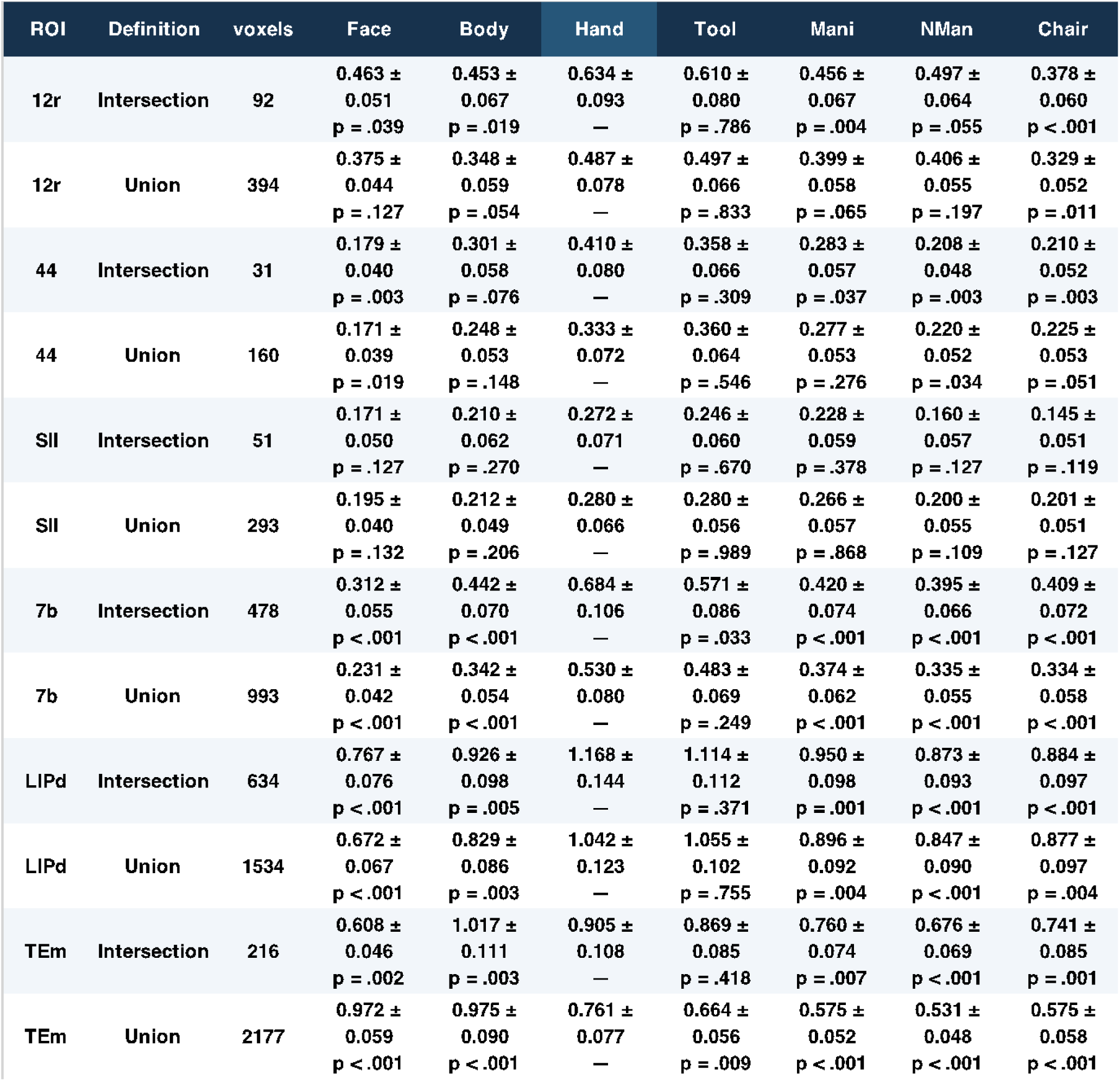
Summary statistics. Summary of the statistical results corresponding to the bar plots in Figure 3, including signal amplitudes and corresponding statistical test (paired t-test, FDR-corrected) results for each fROI and stimulus condition.

**Extended data Table 3.**
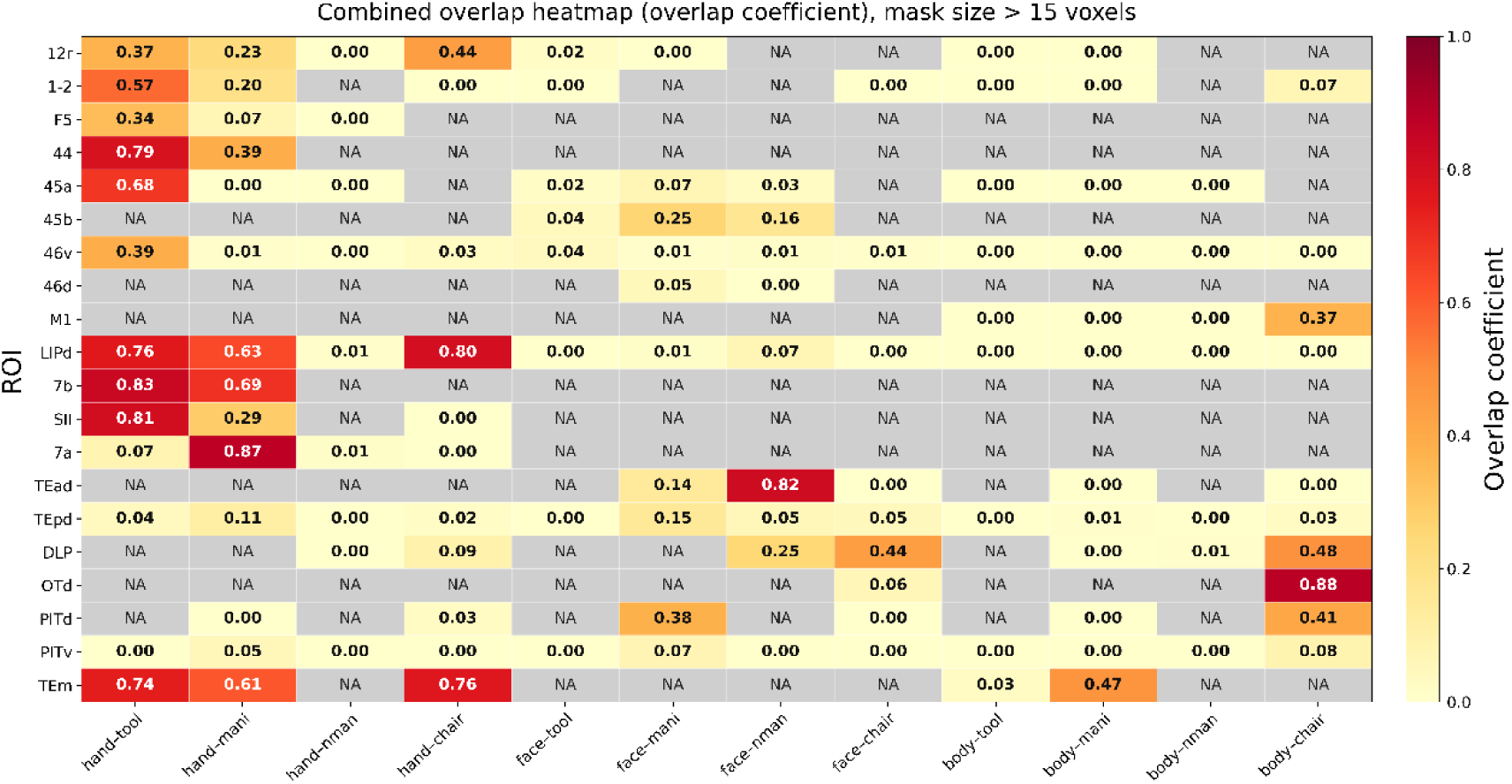
Spatial Overlap Between Animate and Inanimate. Categories summary (for Extended data Figure 5). Spatial overlap coefficients for the same category pairs and atlas-defined cortical areas as in A. Only category pairs in which both corresponding pair-specific functional regions of interest (fROIs) contained more than 15 voxels were included; gray cells marked NA (not applicable) indicate pairs that did not meet this criterion. For each eligible pair, overlap size was defined as the number of voxels belonging to both fROIs, |A ∩ B|, where A and B denote the two pair-specific fROIs. Proportional overlap was quantified using the overlap coefficient, calculated as |A ∩ B| / min(|A|, |B|), where |A| and |B| denote the respective fROI sizes in voxels. This coefficient expresses the proportion of the smaller fROI shared with the larger fROI. Values range from 0 (no spatial overlap) to 1 (complete inclusion of the smaller fROI within the larger). Darker colors indicate greater proportional overlap.

| ROI | hand-tool | hand-mani | hand-nman | hand-chair | face-tool | face-mani | face-nman | face-chair | body-tool | body-mani | body-nman | body-chair |
|---|---|---|---|---|---|---|---|---|---|---|---|---|
| 12r | **0.37** | **0.23** | **0.00** | **0.44** | **0.02** | **0.00** | NA | NA | **0.00** | **0.00** | NA | NA |
| 1-2 | **0.57** | **0.20** | NA | **0.00** | **0.00** | NA | NA | **0.00** | **0.00** | **0.00** | NA | **0.07** |
| F5 | **0.34** | **0.07** | **0.00** | NA | NA | NA | NA | NA | NA | NA | NA | NA |
| 44 | **0.79** | **0.39** | NA | NA | NA | NA | NA | NA | NA | NA | NA | NA |
| 45a | **0.68** | **0.00** | **0.00** | NA | **0.02** | **0.07** | **0.03** | NA | **0.00** | **0.00** | **0.00** | NA |
| 45b | NA | NA | NA | NA | **0.04** | **0.25** | **0.16** | NA | NA | NA | NA | NA |
| 46v | **0.39** | **0.01** | **0.00** | **0.03** | **0.04** | **0.01** | **0.01** | **0.01** | **0.00** | **0.00** | **0.00** | **0.00** |
| 46d | NA | NA | NA | NA | NA | **0.05** | **0.00** | NA | NA | NA | NA | NA |
| M1 | NA | NA | NA | NA | NA | NA | NA | NA | **0.00** | **0.00** | **0.00** | **0.37** |
| LIPd | **0.76** | **0.63** | **0.01** | **0.80** | **0.00** | **0.01** | **0.07** | **0.00** | **0.00** | **0.00** | **0.00** | **0.00** |
| 7b | **0.83** | **0.69** | NA | NA | NA | NA | NA | NA | NA | NA | NA | NA |
| SII | **0.81** | **0.29** | NA | **0.00** | NA | NA | NA | NA | NA | NA | NA | NA |
| 7a | **0.07** | **0.87** | **0.01** | **0.00** | NA | NA | NA | NA | NA | NA | NA | NA |
| TEad | NA | NA | NA | NA | NA | **0.14** | **0.82** | **0.00** | NA | **0.00** | NA | **0.00** |
| TEpd | **0.04** | **0.11** | **0.00** | **0.02** | **0.00** | **0.15** | **0.05** | **0.05** | **0.00** | **0.01** | **0.00** | **0.03** |
| DLP | NA | NA | **0.00** | **0.09** | NA | NA | **0.25** | **0.44** | NA | **0.00** | **0.01** | **0.48** |
| OTd | NA | NA | NA | NA | NA | NA | NA | **0.06** | NA | NA | NA | **0.88** |
| PITd | NA | **0.00** | NA | **0.03** | NA | **0.38** | NA | **0.00** | NA | **0.00** | NA | **0.41** |
| PITv | **0.00** | **0.05** | **0.00** | **0.00** | **0.00** | **0.07** | **0.00** | **0.00** | **0.00** | **0.00** | **0.00** | **0.08** |
| TEm | **0.74** | **0.61** | NA | **0.76** | NA | NA | NA | NA | **0.03** | **0.47** | NA | NA |

**Extended data Table 4.** Statistical summary of the multiple-regression. RSA. Numerical values are shown underlying the regression analysis in Figure 5B. For each fROI and predictor model (CORnet-V1, CORnet-IT, K400 and SSV2), standardized regression coefficients (standardized β) are reported as mean ± S.E.M. across runs. Asterisks indicate regression coefficients significantly different from zero (one-sided sign-flip permutation tests across runs, FDR-adjusted *P* < 0.05) (* < 0.05, ** < 0.01, *** < 0.001).

| ROI | CORnet-V1 | CORnet-IT | K400 | SSV2 |
|---|---|---|---|---|
| 11l | 0.14 ± 0.08 | -0.32 ± 0.10 | -0.24 ± 0.08 | 0.58 ± 0.14** |
| 12r | 0.02 ± 0.10 | -0.10 ± 0.09 | 0.07 ± 0.09 | 0.10 ± 0.10 |
| 44 | -0.11 ± 0.11 | -0.12 ± 0.10 | 0.05 ± 0.05 | 0.18 ± 0.12 |
| 45a | 0.00 ± 0.08 | -0.16 ± 0.08 | 0.38 ± 0.08*** | -0.14 ± 0.07 |
| 46v | -0.29 ± 0.12 | -0.07 ± 0.08 | 0.18 ± 0.08 | 0.21 ± 0.14 |
| F5 | 0.07 ± 0.10 | -0.13 ± 0.11 | -0.08 ± 0.06 | 0.21 ± 0.12 |
| 1-2 | -0.22 ± 0.10 | -0.15 ± 0.08 | -0.05 ± 0.07 | 0.47 ± 0.13** |
| SII | -0.05 ± 0.10 | -0.14 ± 0.11 | 0.04 ± 0.07 | 0.19 ± 0.12 |
| 7a | -0.19 ± 0.10 | -0.32 ± 0.10 | 0.03 ± 0.04 | 0.61 ± 0.13*** |
| 7b | -0.53 ± 0.13 | 0.00 ± 0.09 | 0.11 ± 0.05 | 0.50 ± 0.12*** |
| LIPd | -0.12 ± 0.11 | -0.39 ± 0.10 | 0.05 ± 0.03 | 0.62 ± 0.12*** |
| LIPv | -0.16 ± 0.10 | -0.15 ± 0.08 | 0.24 ± 0.07** | 0.10 ± 0.08 |
| AIP | -0.10 ± 0.10 | -0.23 ± 0.07 | -0.02 ± 0.06 | 0.33 ± 0.11** |
| PITd | 0.59 ± 0.04*** | 0.77 ± 0.12*** | -0.90 ± 0.09 | 0.11 ± 0.07 |
| PITv | 0.54 ± 0.07*** | -0.04 ± 0.12 | -0.54 ± 0.11 | 0.44 ± 0.08*** |
| TEa | -0.45 ± 0.08 | 0.96 ± 0.17*** | -0.55 ± 0.09 | 0.16 ± 0.06* |
| TEm | 0.31 ± 0.06*** | 0.63 ± 0.19** | -0.60 ± 0.11 | 0.16 ± 0.07* |
| TEpd | 0.27 ± 0.07*** | 0.11 ± 0.13 | -0.19 ± 0.10 | 0.10 ± 0.09 |
| DLP | 0.60 ± 0.07*** | 0.15 ± 0.14 | -0.43 ± 0.09 | 0.13 ± 0.08 |
| V4 | 0.58 ± 0.07*** | 0.21 ± 0.13 | -0.51 ± 0.09 | 0.07 ± 0.08 |
| V3d | 0.39 ± 0.08*** | 0.22 ± 0.11 | -0.29 ± 0.12 | 0.06 ± 0.08 |
| V3a | 0.55 ± 0.07*** | 0.43 ± 0.11** | -0.48 ± 0.11 | -0.06 ± 0.05 |
| V2v | 0.01 ± 0.06 | -0.27 ± 0.07 | 0.43 ± 0.08*** | 0.15 ± 0.07 |
| V2d | 0.47 ± 0.06*** | 0.50 ± 0.08*** | -0.12 ± 0.10 | -0.20 ± 0.05 |
| V1 | 0.70 ± 0.05*** | 0.18 ± 0.11 | -0.36 ± 0.09 | -0.07 ± 0.06 |
| 8Bs | 0.22 ± 0.09* | -0.25 ± 0.08 | -0.09 ± 0.10 | 0.03 ± 0.07 |
| FST | 0.20 ± 0.06** | 0.62 ± 0.13*** | -0.30 ± 0.12 | -0.13 ± 0.07 |
| LOP | 0.00 ± 0.07 | 0.63 ± 0.17** | -0.38 ± 0.13 | -0.21 ± 0.06 |
| OTd | 0.67 ± 0.04*** | -0.35 ± 0.07 | -0.39 ± 0.07 | 0.69 ± 0.06*** |

## Notes

### Competing Interest Statement

The authors have declared no competing interest.

## References

1. Grill-Spector, K. & Weiner, K. S. The functional architecture of the ventral temporal cortex and its role in categorization. Nature Reviews Neuroscience vol. 15 Preprint at 10.1038/nrn3747 (2014).

2. Konkle, T. & Caramazza, A. Tripartite organization of the ventral stream by animacy and object size. Journal of Neuroscience 33, (2013).

3. Bell, A. H., Hadj-Bouziane, F., Frihauf, J. B., Tootell, R. B. H. & Ungerleider, L. G. Object representations in the temporal cortex of monkeys and humans as revealed by functional magnetic resonance imaging. J. Neurophysiol. 101, (2009).

4. Bao, P., She, L., McGill, M. & Tsao, D. Y. A map of object space in primate inferotemporal cortex. Nature 583, (2020).

5. Kriegeskorte, N. et al. Matching Categorical Object Representations in Inferior Temporal Cortex of Man and Monkey. Neuron 60, (2008).

6. Chao, L. L. & Martin, A. Representation of manipulable man-made objects in the dorsal stream. Neuroimage 12, (2000).

7. Johnson-Frey, S. H. et al. Actions or hand-object interactions? Human inferior frontal cortex and action observation. Neuron 39, (2003).

8. Lewis, J. W. Cortical networks related to human use of tools. Neuroscientist vol. 12 Preprint at 10.1177/1073858406288327 (2006).

9. Bracci, S., Caramazza, A. & Peelen, M. V. View-invariant representation of hand postures in the human lateral occipitotemporal cortex. Neuroimage 181, (2018).

10. Bracci, S., Cavina-Pratesi, C., Ietswaart, M., Caramazza, A. & Peelen, M. V. Closely overlapping responses to tools and hands in left lateral occipitotemporal cortex. J. Neurophysiol. 107, (2012).

11. Cortinovis, D., Truong, N., Op de Beeck, H. & Bracci, S. Investigating action topography in visual cortex and deep artificial neural networks. Nature Communications 17, (2026).

12. Bracci, S., Daniels, N. & De Beeck, H. O. Task context overrules object-And category-related representational content in the human parietal cortex. Cerebral Cortex 27, (2017).

13. Mahon, B. Z. & Almeida, J. Reciprocal interactions among parietal and occipito-temporal representations support everyday object-directed actions. Neuropsychologia 198, (2024).

14. Orban, G. A., Van Essen, D. & Vanduffel, W. Comparative mapping of higher visual areas in monkeys and humans. Trends in Cognitive Sciences vol. 8 Preprint at 10.1016/j.tics.2004.05.009 (2004).

15. Vanduffel, W., Zhu, Q. & Orban, G. A. Monkey Cortex through fMRI Glasses. Neuron vol. 83 Preprint at 10.1016/j.neuron.2014.07.015 (2014).

16. Peeters, R. et al. The representation of tool use in humans and monkeys: Common and uniquely human features. Journal of Neuroscience 29, (2009).

17. Iriki, A., Tanaka, M. & Iwamura, Y. Coding of modified body schema during tool use by macaque postcentral neurones. Neuroreport 7, (1996).

18. Perrett, D. I. et al. Visual analysis of body movements by neurones in the temporal cortex of the macaque monkey: A preliminary report. Behavioural Brain Research 16, (1985).

19. Jellema, T., Baker, C. I., Wicker, B. & Perrett, D. I. Neural representation for the perception of the intentionality of actions. Brain Cogn. 44, (2000).

20. Fogassi, L., et al. Neuroscience: Parietal lobe: From action organization to intention understanding. Science (1979). 308, (2005).

21. Nelissen, K. & Vanduffel, W. Grasping-related functional magnetic resonance imaging brain responses in the macaque monkey. Journal of Neuroscience 31, (2011).

22. Nelissen, K. et al. Action observation circuits in the macaque monkey cortex. Journal of Neuroscience 31, (2011).

23. Pani, P., Theys, T., Romero, M. C. & Janssen, P. Grasping execution and grasping observation activity of single neurons in the macaque anterior intraparietal area. J. Cogn. Neurosci. 26, (2014).

24. Li, X., Zhu, Q. & Vanduffel, W. Submillimeter fMRI reveals an extensive, fine-grained and functionally-relevant scene-processing network in monkeys. Prog. Neurobiol. 211, (2022).

25. Janssens, T. et al. An implanted 8-channel array coil for high-resolution macaque MRI at 3T. Neuroimage 62, (2012).

26. Fitzgerald, N. E., Montaldo, G., Froesel, M., Urban, A. & Vanduffel, W. Volumetric Functional Ultrasound Imaging in Macaques. IEEE Trans. Med. Imaging 45, (2026).

27. Balan, P. F. et al. MEBRAINS 1.0: A new population-based macaque atlas. Imaging Neuroscience 2, (2024).

28. Fitzgerald Nora E., Zhu Qi, Palomero-Gallagher Nicola & Vanduffel Wim. MEBRAINS: a new population-based monkey template (v3.0). Preprint at 10.25493/P7JR-DRG (2026).

29. Chao, L. L., Haxby, J. V. & Martin, A. Attribute-based neural substrates in temporal cortex for perceiving and knowing about objects. Nat. Neurosci. 2, (1999).

30. Sha, L. et al. The animacy continuum in the human ventral vision pathway. J. Cogn. Neurosci. 27, (2015).

31. Popivanov, I. D., Jastorff, J., Vanduffel, W. & Vogels, R. Stimulus representations in body-selective regions of the macaque cortex assessed with event-related fMRI. Neuroimage 63, (2012).

32. Tsao, D. Y., Freiwald, W. A., Knutsen, T. A., Mandeville, J. B. & Tootell, R. B. H. Faces and objects in macaque cerebral cortex. Nat. Neurosci. 6, (2003).

33. Pinsk, M. A., DeSimone, K., Moore, T., Gross, C. G. & Kastner, S. Representations of faces and body parts in macaque temporal cortex: A functional MRI study. Proc. Natl. Acad. Sci. U. S. A. 102, (2005).

34. Mergan, E., Zhu, Q., Li, X., Vogels, R. & Vanduffel, W. Fast face-selective responses in prefrontal face patches of the macaque. Cell Rep. 44, (2025).

35. Kubilius, J. et al. CORnet: Modeling the Neural Mechanisms of Core Object Recognition. bioRxiv Preprint at 10.1101/408385 (2018).

36. Kubilius, J. et al. Brain-like object recognition with high-performing shallow recurrent ANNs. in Advances in Neural Information Processing Systems vol. 32 (2019).

37. Will Kay etc. The Kinetics Human Action Video Dataset. International Journal of Mathematical Analysis (2009).

38. Goyal, R. et al. The ‘Something Something’ Video Database for Learning and Evaluating Visual Common Sense. in Proceedings of the IEEE International Conference on Computer Vision vols 2017-October (2017).

39. Liu, Z. et al. Video Swin Transformer. in Proceedings of the IEEE Computer Society Conference on Computer Vision and Pattern Recognition vols 2022-June (2022).

40. Mumford, J. A., Poline, J. B. & Poldrack, R. A. Orthogonalization of regressors in fMRI models. PLoS One 10, (2015).

41. Cometa, A. et al. Brain and grammar: revealing electrophysiological basic structures with competing statistical models. Cerebral Cortex 34, (2024).

42. Bracci, S. & Peelen, M. V. Body and object effectors: The organization of object representations in high-level visual cortex reflects body-object interactions. Journal of Neuroscience 33, (2013).

43. Bracci, S., Caramazza, A. & Peelen, M. V. Representational similarity of body parts in human occipitotemporal cortex. Journal of Neuroscience 35, (2015).

44. Bracci, S., Ietswaart, M., Peelen, M. V. & Cavina-Pratesi, C. Dissociable neural responses to hands and non-hand body parts in human left extrastriate visual cortex. J. Neurophysiol. 103, (2010).

45. Vaesen, K. The cognitive bases of human tool use. Behavioral and Brain Sciences 35, (2012).

46. Lesourd, M. et al. Semantic and action tool knowledge in the brain: Identifying common and distinct networks. Neuropsychologia vol. 159 Preprint at 10.1016/j.neuropsychologia.2021.107918 (2021).

47. Peelen, M. V. et al. Tool selectivity in left occipitotemporal cortex develops without vision. J. Cogn. Neurosci. 25, (2013).

48. Dehaene, S. & Cohen, L. Cultural recycling of cortical maps. Neuron vol. 56 Preprint at 10.1016/j.neuron.2007.10.004 (2007).

49. Jellema, T. & Perrett, D. I. Cells in monkey STS responsive to articulated body motions and consequent static posture: A case of implied motion? Neuropsychologia 41, (2003).

50. Zafirova, Y., Bahrami, A. & Vogels, R. Body-part tuning in the macaque superior temporal sulcus. Cerebral Cortex 36, (2026).

51. Messinger, A., Squire, L. R., Zola, S. M. & Albright, T. D. Neuronal representations of stimulus associations develop in the temporal lobe during learning. Proc. Natl. Acad. Sci. U. S. A. 98, (2001).

52. Li, N. & DiCarlo, J. J. Unsupervised natural visual experience rapidly reshapes size-invariant object representation in inferior temporal cortex. Neuron 67, (2010).

53. Meyer, T. & Olson, C. R. Statistical learning of visual transitions in monkey inferotemporal cortex. Proc. Natl. Acad. Sci. U. S. A. 108, (2011).

54. Woloszyn, L. & Sheinberg, D. L. Effects of Long-Term Visual Experience on Responses of Distinct Classes of Single Units in Inferior Temporal Cortex. Neuron 74, (2012).

## Methods reference

55. Vanduffel, W. et al. Visual motion processing investigated using contrast agent-enhanced fMRI in awake behaving monkeys. Neuron 32, (2001).

56. Zhu, Q. & Vanduffel, W. Submillimeter fMRI reveals a layout of dorsal visual cortex in macaques, remarkably similar to New World monkeys. Proc. Natl. Acad. Sci. U. S. A. 116, (2019).

57. Willenbockel, V. et al. Controlling low-level image properties: The SHINE toolbox. Behav. Res. Methods 42, (2010).

58. Matić, K., Op de Beeck, H. & Bracci, S. It’s not all about looks: The role of object shape in parietal representations of manual tools. Cortex 133, (2020).

59. Avants, B. B. et al. A reproducible evaluation of ANTs similarity metric performance in brain image registration. Neuroimage 54, (2011).

60. Fedorov, A. et al. 3D Slicer as an image computing platform for the Quantitative Imaging Network. Magn. Reson. Imaging 30, (2012).

61. Wang, Z., Bovik, A. C., Sheikh, H. R. & Simoncelli, E. P. Image quality assessment: From error visibility to structural similarity. IEEE Transactions on Image Processing 13, (2004).

62. Li, X., Zhu, Q., Janssens, T., Arsenault, J. T. & Vanduffel, W. In Vivo Identification of Thick, Thin, and Pale Stripes of Macaque Area V2 Using Submillimeter Resolution (f)MRI at 3 T. Cerebral Cortex 29, (2019).

63. Benjamini, Yoav; Hochberg, Y. Controlling the False Discovery Rate - a Practical and Powerful Approach to Multiple Testing. Journal of the Royal Statistical Society Series B-Methodological 1995.pdf. Journal of the Royal Statistical Society Series B (Methological) vol. 57 Preprint at (1995).

64. Walther, A. et al. Reliability of dissimilarity measures for multi-voxel pattern analysis. Neuroimage 137, (2016).

65. Diedrichsen, J. & Kriegeskorte, N. Representational models: A common framework for understanding encoding, pattern-component, and representational-similarity analysis. PLoS Comput. Biol. 13, (2017).

66. Pelekanos, V. et al. Rapid event-related, BOLD fMRI, non-human primates (NHP): choose two out of three. Sci. Rep. 10, (2020).

67. Winkler, A. M., Ridgway, G. R., Webster, M. A., Smith, S. M. & Nichols, T. E. Permutation inference for the general linear model. Neuroimage 92, (2014).

